# Sepsis Induces Age- and Sex-Specific Chromatin Remodeling in Myeloid-Derived Suppressor Cells

**DOI:** 10.1101/2025.05.26.656229

**Authors:** Angel M. Charles, Christine E. Rodhouse, Dijoia B. Darden, Marie-Pierre L. Gauthier, Mingqi Zhou, Miguel Hernandez-Rios, Dayuan Wang, Gemma Casadesus, Letitia Bible, Alicia M. Mohr, Feifei Xiao, Guoshuai Cai, Jason O. Brant, Shannon M. Wallet, Clayton E. Mathews, Lyle L. Moldawer, Paramita Chakrabarty, Rhonda L. Bacher, Philip A. Efron, Robert Maile, Michael P. Kladde

## Abstract

Sepsis survivors frequently develop long-term immune dysfunction, but the epigenetic mechanisms underlying persistent myeloid suppression remain unclear. Myeloid-derived suppressor cells (MDSCs), whose function is shaped by host age and sex, are key contributors to post-sepsis immune dysregulation. Here, we present a high-resolution epigenetic map targeting gene promoters of MDSCs after sepsis using MAPit-FENGC, a single-molecule assay that simultaneously profiles DNA methylation and chromatin accessibility. In a clinically relevant murine model including young and older adult male and female mice, splenic MDSCs were isolated for MAPit-FENGC and single-cell RNA sequencing. Unsupervised clustering identified nine promoter classes reflecting chromatin dynamics: age- and sex-dependent sepsis-induced opening (Classes 1-4), persistent closure with varying levels of DNA methylation (Classes 5-7), and constitutive openness post-sepsis (Classes 8, 9). Transcriptomic profiling corroborated these promoter states, linking accessibility with gene expression. These findings establish how epigenetic reprogramming of MDSCs may shape age- and sex-specific immune trajectories in sepsis survivors.

## INTRODUCTION

Sepsis, when the host has organ insufficiency/failure in response to infection^1^ affects in excess of 1.7 million adults in the United States, and results in the death of more than 350,000 of these individuals.^2^ In addition, sepsis is estimated to cost greater than $38 billion per year in America alone.^3^ Although acute hospital mortality has improved due to measures such as the Surviving Sepsis campaign,^4^ long-term patient outcomes remain dismal.^5,6^ Despite the significance and impact of all the former, few to no therapeutics are available for this patient population.^7^

Myeloid-derived suppressor cells (MDSCs), delineated almost two decades ago,^8^ are defined as pathologically activated myeloid cells with potent immunosuppressive activity, yet they also possess some pro-inflammatory properties.^9^ Although the leukocytes have been investigated in the cancer field for some time, research focused on MDSCs in sepsis is relatively recent.^10^ MDSCs have been demonstrated to expand in number, remain circulating in the sepsis survivor, and are associated with poor outcomes in these patients.^11,12^ Whereas these sepsis-derived MDSCs meet the key functional definition of MDSCs (immunosuppressive activity against lymphocytes, etc.), data indicate that MDSCs from septic mice are not identical to oncologic-derived MDSCs, in both mice and humans.^13–15^ Modification of MDSCs in the sepsis survivor represents a potential approach to improve these patients’ outcomes,^16^ but still requires a better disease-specific understanding of these cells.

In addition to the need for more disease-specific precision/personalized therapeutics for sepsis, there is a need to delineate how sepsis engenders unique effects in certain cohorts, e.g., age and sex.^16–25^ Work from our group and others has illustrated that both sex and age have significant effects on the sepsis response, as well as on the post-infection MDSCs generated by that host.^17,18,20–22^ Thus, investigating these cohorts will be key to any successful future interventions for sepsis survivors.

Epigenetics, the study of alterations in gene expression not due to changes in the genetic code, are considered a potential method of successfully modifying MDSCs in the host so that these cells will no longer be pathologic.^26^ One such method of studying the epigenetics of the sepsis-induced MDSC is Methyltransferase Accessibility Protocol for Individual Templates combined with Flap-Enabled Next-Generation Capture (MAPit-FENGC), a validated method that allows for simultaneous single-molecular-level analysis of endogenous CpG methylation and accessibility at targeted promoter and enhancer regions.^27^ Using a preclinical surgical sepsis animal model that better parallels the human condition,^17,21^ we utilized MAPit to assess the epigenetic uniqueness of isolated murine splenic Cd11b^+^Gr1^+^ MDSCs across several cohorts: sepsis versus control, male versus female, and young adult versus older adult.

## RESULTS

Sepsis exerted a profound and consistent remodeling of the MDSC epigenome. Promoters that regulate inflammatory, metabolic, and differentiation-associated programs displayed striking shifts in both accessibility and methylation patterns. Specifically, multiple gene promoters showed increased accessibility paired with reduced methylation, a configuration associated with transcriptional activation, while other loci demonstrated persistent closure and hypermethylation, consistent with durable repression. This dichotomy defined nine distinct promoter classes, of which Classes 1-4 reflected sepsis-induced opening, whereas Classes 5-7 and Classes 8, 9 captured states with persistently closed (likely repressed) and constitutively open (likely expressed) chromatin, respectively. Importantly, these global patterns were evident regardless of host age or sex, underscoring sepsis itself as the primary driver of epigenetic reprogramming.

### Global epigenetic structure separates MDSCs by sepsis, age, and sex

Principal Component Analysis (PCA) of MAPit-FENGC single-molecule data revealed clear global distinctions (out to four PCs) in the epigenetic landscape of CD11b^+^Gr1⁺ splenic MDSCs based on sepsis exposure, age, and sex (**Figures S1, S2**). In PCA of endogenous CpG methylation alone (**Figure S1**), septic mice formed distinct clusters compared to naïve controls. Notably, older adult females demonstrated the greatest divergence in CpG methylation patterns after sepsis, suggesting age- and sex-dependent remodeling of the methylome. When PCA was restricted to GpC methylation (chromatin accessibility) data (**Figure S2**), sepsis again drove marked separation, particularly among female samples. Septic females (both young and older adult) clustered distinctly from their naïve counterparts, whereas MDSCs derived from all adult males displayed a more intermediate shift. This suggests that chromatin accessibility is broadly altered by sepsis, with a stronger response in females.

### Distinct promoter accessibility classes exist in MDSCs following sepsis

Unsupervised clustering analysis^28^ of individual MAPit-FENGC promoter copies revealed nine promoter classes (Classes 1-9) in splenic CD11b^+^Gr1^+^ MDSCs, each with characteristic chromatin accessibility and DNA methylation patterns across conditions (**Table S1**). These classes captured the range of epigenetic responses to sepsis and underline strong age- and sex-specific differences in MDSC promoter regulation. Here, we describe qualitative trends for each class, highlighting how sepsis-induced chromatin accessibility (nucleosome-free regions, NFRs) and DNA methylation changes vary by promoter class, age, and sex.

#### 1. Sepsis induces NFRs at many promoters (Classes 1-4)

Several promoter classes exhibited marked chromatin opening due to nucleosome sliding/eviction in response to sepsis. A single Class 1 promoter, *S100a9* (encoding the calcium-binding protein A9, a key MDSC protein that regulates MDSC trafficking, expansion, and activation, as well as acting as an alarmin),^29,30^ showed dramatic, sepsis-driven increases in accessibility (at GCH sites; each mouse, right within each pair of panels; yellow color; **Figure 1A**). Large NFRs, with peak accessibility upstream of the transcription start site (TSS), formed on *S100a9* promoter copies in five of six total septic female (**Figure 1A**) and male (not shown) mice. These NFRs were characterized by promoter copies with continuous spans of accessible GCH of variable length— some extending across the entire 523 bp assayed region. The magnitude of *S100a9* NFR formation in Class 1 was modulated by sex and age. In MDSCs isolated from females, older septic mice had significantly more extensive *S100a9* promoter opening than young septic females, whereas male MDSCs showed robust *S100a9* NFR formation in sepsis regardless of age (though slightly less pronounced than in females). In septic mice, *S100a9* demonstrated increased average chromatin accessibility in older adult females compared to young adult females and older adult males, with older females showing the largest NFRs and greatest fraction of molecules with NFRs (indicating a higher proportion of cells with an open promoter; **Figure 1B**). This was accompanied by a modest decrease in average CpG (at HCGs) methylation across the locus, most notably in the proximal promoter region. This Class 1 promoter had moderate DNA methylation at baseline that underwent partial demethylation upon sepsis, in line with an activated chromatin state (**Figure 1C)**.

**Figure 1.**
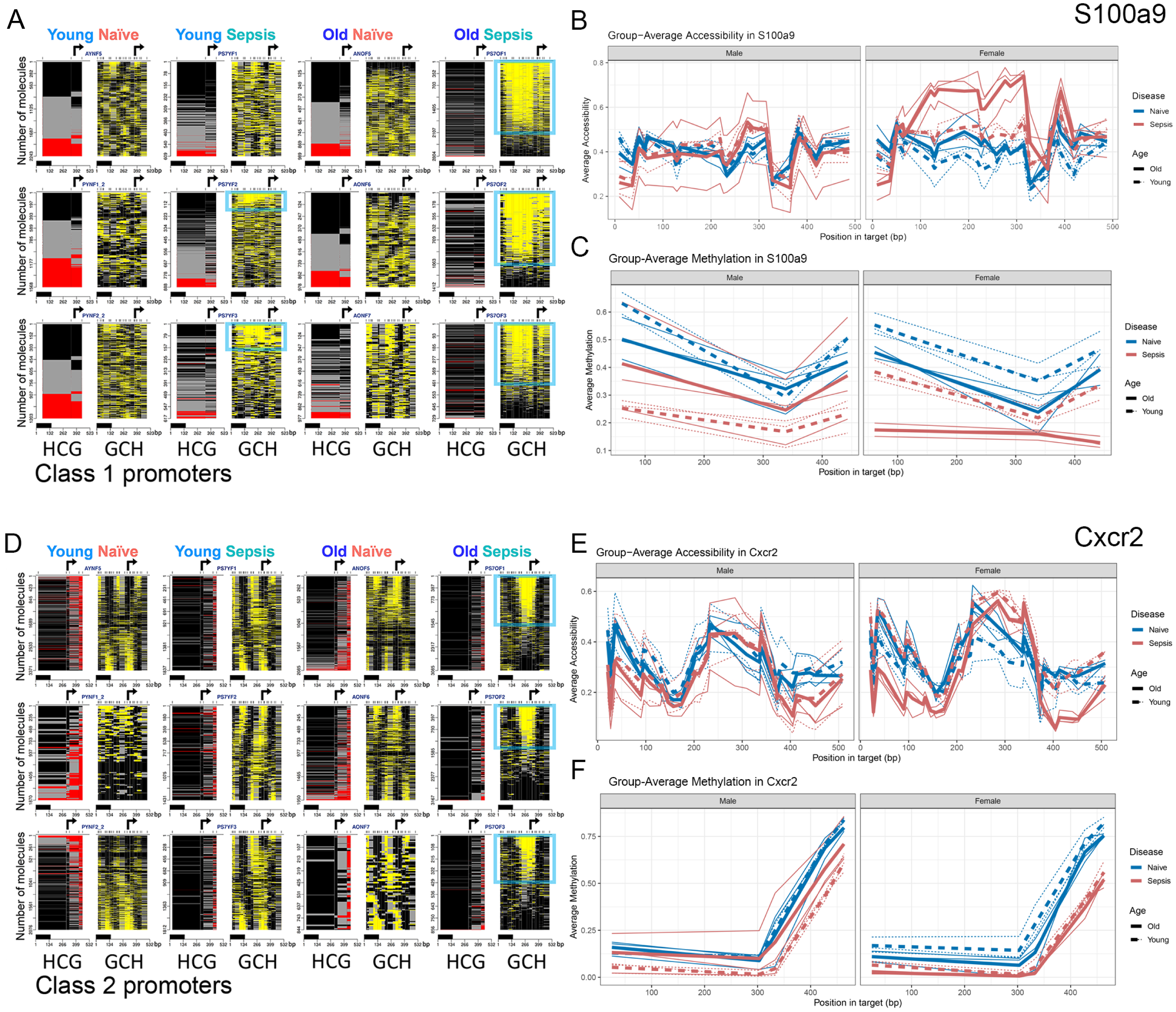
Class 1 and 2 promoters exhibit robust sepsis-induced chromatin accessibility, with age- and sex-specific differences and partial demethylation across MDSC subsets (A) Representative single-molecule MAPit-FENGC plots for selected Class 1 promoters, showing simultaneous profiling of endogenous CpG methylation (HCG, left of each pair of panels, red) and *in vitro* GpC methylation (GCH, right of each pair, yellow) in Gr1⁺ MDSCs from four experimental groups: Young Naïve, Young Sepsis, Older Naïve, and Older Sepsis, all from female mice. Each horizontal line represents a unique DNA molecule, with the total number indicated on the y-axis. Black and gray marks denote methylated cytosines, while yellow marks indicate unmethylated (open or accessible) cytosines. Promoters are organized by gene and condition; Class 1 promoters were defined by consistent nucleosome-free region (NFR) formation (highlighted by blue boxes) in all septic groups, which co-localizes with the TSS (arrow). The black bar on top of the base pair scale is the size of a nucleosome particle (147 bp). Particularly robust NFRs are observed in *Older Sepsis* conditions, especially in females. (B) Group-averaged chromatin accessibility (GpC methylation) across the same *S100a9* region, plotted separately for males (left) and females (right). (C) Group-averaged endogenous CpG methylation levels across the *S100a9* promoter. (D-F) Equivalent plots for the Class 2 promoter genes *Cxcr2* in Gr1⁺ MDSCs.

Class 2 promoters also gained accessibility in septic conditions but in a more context-dependent manner. Specifically, Class 2 chromatin opened (NFRs appeared) in older adult female sepsis and in male sepsis (both young and older), but not in young female sepsis. This pattern indicates that young females failed to open these promoters unless aged, whereas males of either age mounted an increase in accessibility. When present, the degree of NFR formation for Class 2 genes was similar between sexes. A representative Class 2 gene is *Cxcr2* (**Figure 1D**; encodes C-X-C chemokine receptor type 2, trafficking and facilitating the suppressive function of MDSCs within inflamed tissue).^31^ In MDSCs from septic hosts, the *Cxcr2* promoter remained nucleosome-occupied (by randomly positioned nucleosomes) in young females but showed clear NFR formation at the TSS in older females (and in males of both ages). Concordantly, *Cxcr2* and other Class 2 promoters (e.g., *Nos2*, *Ptgs2*) exhibited low-to-moderate DNA methylation (at HCGs; left; red color) that either remained stable or slightly decreased with sepsis in the responsive groups (**Figure 1D**). The requirement of advanced age for female MDSCs to demonstrate open Class 2 loci, *versus* no such requirement in males, highlights a female-specific age dependency in epigenetic activation of this promoter set. Average chromatin accessibility at the *Cxcr2* locus was significantly increased in septic older adult mice, particularly females, consistent with an epigenetic state favoring enhanced expression (**Figure 1E**). This accessibility gain was not accompanied by substantial changes in DNA methylation (**Figure 1F**), which remained uniformly low upstream of the TSS across groups, suggesting a permissive chromatin environment at baseline, with further inducibility following septic challenge.

Class 3 promoters were broadly responsive to sepsis across all cohorts, with no significant sex or age differences in their accessibility changes. These loci, e.g., *S100a8* (S100A8 acts similarly to S100A9^29^) and *Mmp8* (MMP8 facilitates MDSC mobilization and migration, as well as the immunosuppressive environment),^32^ formed NFRs in MDSCs from septic hosts in both males and females, and in both young and older mice, to a similar extent. As a result, Class 3 represents a core set of promoters that are consistently opened by sepsis irrespective of host age or sex. DNA methylation of Class 3 promoters was generally low to intermediate and showed modest changes (often slight demethylation), if any, upon sepsis. The uniform NFR formation at these sites suggests they may drive fundamental sepsis-responsive genes in MDSCs that are required in all populations. For instance, *S100a8* (encodes the S100 calcium-binding protein A8) promoter exhibited equivalent accessibility gains in septic males and females, young and older (data not shown), mirroring the behavior of its Class 1 counterpart *S100a9*, but without the female-older bias. Similarly, *Mmp8* (encoding matrix metalloproteinase-8) opened consistently in all septic groups, forming an NFR over the TSS (**Figure 2A**). Thus, Class 3 promoters highlight sepsis-induced epigenetic changes that are common to both sexes and ages, representing an invariant component of the MDSC response to sepsis. *Mmp8* showed a sepsis-induced increase in chromatin accessibility (**Figure 2B**) alongside a pronounced decrease in DNA methylation in its promoter region (**Figure 2C**). These coordinated epigenetic modifications suggest enhanced transcriptional readiness.

**Figure 2.**
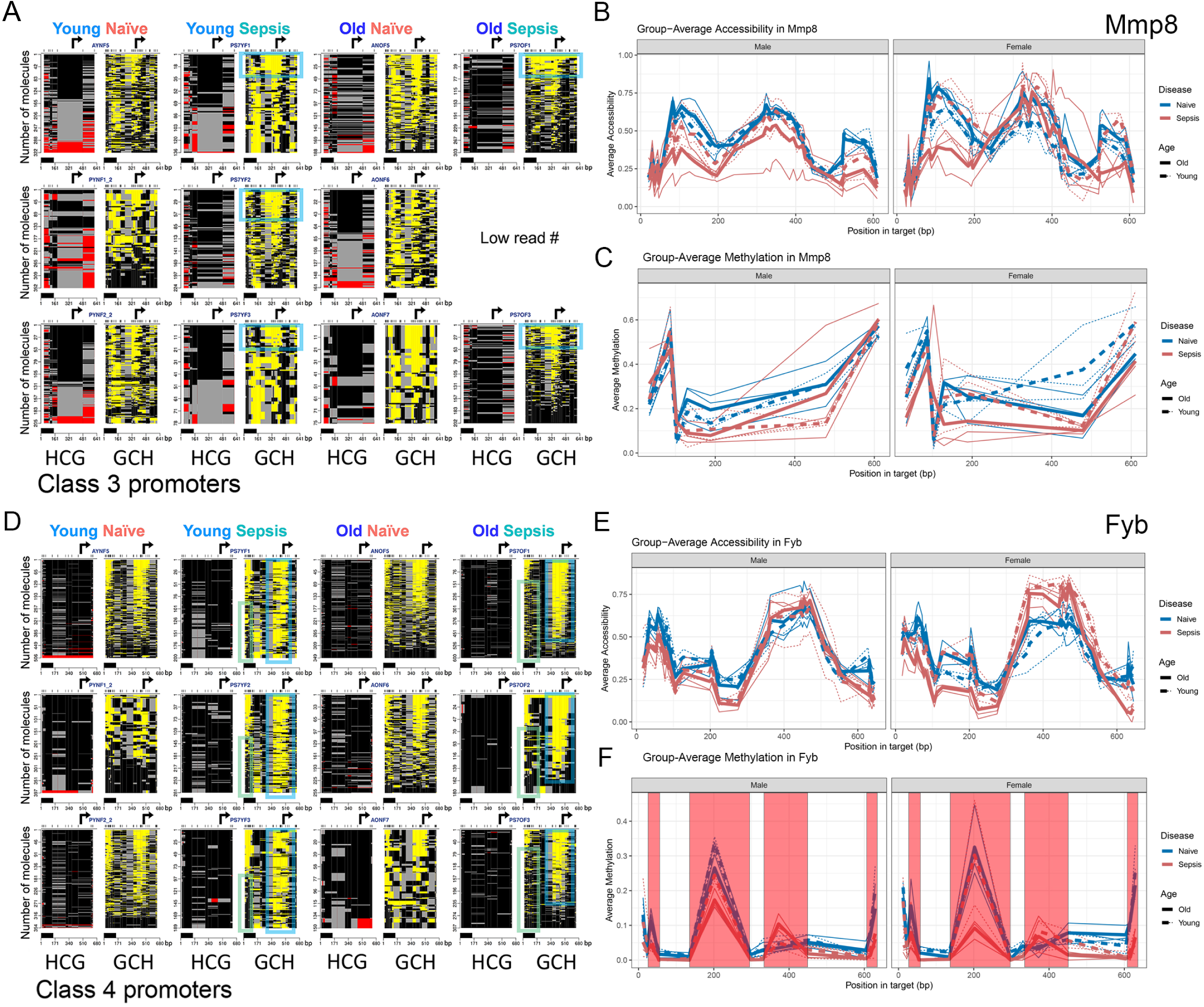
Class 3 promoters are consistently accessible in sepsis across all age and sex groups, and Class 4 promoters are activated by sepsis in females only (A) Single-molecule MAPit-FENGC plots for Class 3 genes (e.g., *Mmp8*), showing HCG (left) and GCH (right) signals across Young/Older Female MDSCs. Sepsis induces NFRs in all septic conditions, regardless of age. The figure is annotated as in Figure 1. (B) Group-averaged chromatin accessibility (GpC methylation) across the same region, plotted separately for males (left) and females (right). (C) Group-averaged endogenous CpG methylation levels across the promoter. (D-F) Equivalent plots for the Class 4 promoter *Fyb.* The additional green boxes in (D) highlight areas of protection against the GpC DNA methyltransferase chromatin probe, which accompanies NFR formation in MDSCs from female septic mice. Red shading in (F) indicates sequences that contain CCG sites, an off-target specificity of the GpC enzyme, which therefore do not reflect *bona fide* CpG methylation.

In contrast, a single Class 4 promoter displayed a clear sex-specific accessibility pattern. This promoter, *Fyb*, drives expression of adhesion and degranulation adaptor protein (ADAP), a signaling adaptive protein hypothesized to support MDSC migration, suppressive activity, and survival.^33^ *Fyb* promoter demonstrated TSS-localized NFRs in older and young adult naïve mice (**Figure 2D**), and an increased number of NFR-containing MDSCs from septic female mice of both ages. Concomitantly, in septic mice, an upstream region of accessibility in naïve mice became inaccessible in a subset of *Fyb* promoter molecules after septic shock (**Figure 2D**; green rectangle). In contrast, septic males (older or young) showed no detectable increase in chromatin opening or the concomitant protection beyond the initial accessibility of the *Fyb* TSS region in naïve male mice of both ages. The average chromatin accessibility plot reveals increased and decreased accessibility in MDSCs from septic mice compared to naïve controls over the TSS region and upstream regulatory elements of *Fyb*, respectively (**Figure 2E**). Complementary to this, the average methylation track shows DNA methylation levels at the Class 4 *Fyb* promoter were rather low at baseline and showed further demethylation in female sepsis, especially in older mice, consistent with an epigenetic landscape that supports transcriptional activation in females (**Figure 2F**). The absence of any chromatin change in males, despite an adequate sepsis insult (as verified by other classes), emphasizes a sex divergence in epigenetic regulation, with preferential activation of the Class 4 locus in MDSCs from female mice during sepsis.

#### 2. Persistently closed promoters remain refractory to induction in sepsis (Classes 5-7)

In contrast to the sepsis-inducible classes above, chromatin at Class 5-7 loci was closed with no formation of an NFR in any group (naïve or septic, male or female, young or older). Instead, accessibility was limited to relatively short linkers distributed across each promoter, indicating that their DNA was packaged into arrays of randomly positioned nucleosomes. Notably, linker accessibility in Class 5-7 promoters decreased in MDSCs isolated from septic mice (older females and males) and young septic male mice. Therefore, Classes 5-7 represent promoters that underwent sepsis-specific loss of linker accessibility in an age- and sex-driven manner, contrasting with the NFR formation observed in Class 1-4 promoters.

The distinction between Classes 5, 6, and 7 lies in their DNA methylation status: Class 5 promoters were generally characterized by high levels of endogenous DNA methylation at all CpGs (hypermethylation), which is consistent with a silenced state; Class 6 promoters exhibited moderate-to-high CpG methylation (typically showing little to no demethylation); and Class 7 promoters had low baseline methylation. Despite these differences, all three classes demonstrated a sepsis-specific loss of accessibility in older mice (male and female), and to a lesser extent, in young adult male mice.

Representative Class 5 genes include *Lyz1* (lysozyme 1), *Retn* (resistin), and *F7* (coagulation factor VII), among others (**Figure 3A-C**). After sepsis, these promoters exhibited minimal change in DNA methylation but showed decreased linker accessibility, specifically, in older adult mice (both sexes) and in young males. For example, *Lyz1* promoter remained highly methylated, with no visible formation of large NFRs capable of supporting active transcription in MDSCs from either naive or septic mice. Similarly, *Retn* showed no NFR formation in any condition, although its methylation level declined slightly in older sepsis compared to older naïve animals.

**Figure 3.**
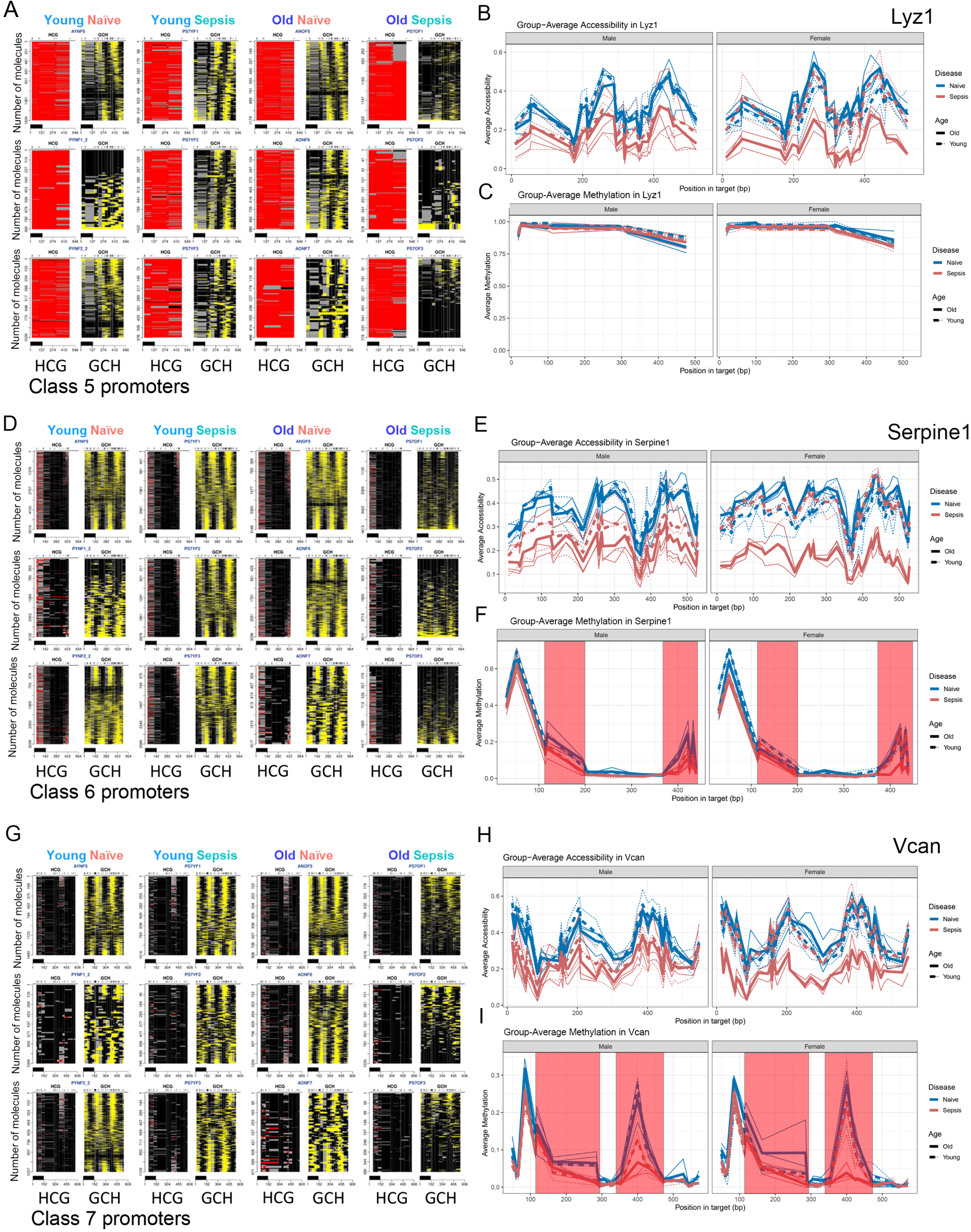
Class 5-7 promoters possess varying levels of DNA methylation but are all packaged into arrays of positioned nucleosomes, where linkers lose accessibility in Older Sepsis only (A) MAPit-FENGC plots of *Lyz1* showing no detectable NFRs in any condition across MDSCs from young/older female mice. HCG and GCH signals indicate high methylation and random nucleosome positioning. The figure is annotated as in Figures 1 and 2. (B) Group-averaged chromatin accessibility (GpC methylation) across the same region, plotted separately for males (left) and females (right). (C) Group-averaged endogenous CpG methylation levels across the promoter. (D-F) Equivalent plots for the Class 6 *Serpine1* promoter. (G-I) Equivalent plots for the Class 7 *Vcan* promoter.

Exemplifying Class 6, the *Serpine1* promoter, encoding the fibrinolytic serine protease inhibitor Pai-1, a protein that influences the presence and activity of MDSCs, often promoting immunosuppression and tumor growth)^34^ was occupied by arrays of randomly positioned nucleosomes in naïve control mice (**Figure 3D-F**). Similar to Class 5 promoters, much of this characteristic linker accessibility was diminished in MDSCs from older septic mice (**Figure 3D**). Average chromatin accessibility plots show that *Serpine1* is less accessible in septic groups, particularly in older mice, and to a lesser extent, young males, compared to their naïve counterparts (**Figure 3E**). No true DNA demethylation was detected (**Figure 3F**; the apparent demethylation of five CpG sites around map position 423 bp in MDSCs from some septic mice is due to off-target methylation by M.CviPI at accessible CCG sites (region highlighted red), and therefore instead reflects loss of chromatin accessibility at those CCGs). Despite baseline CpG methylation at the Serpine 1 TSS (map position 367 bp), the chromatin state of the *Serpine1* promoter is consistent with strong transcriptional silencing due to occupancy by randomly positioned nucleosomes and absence of NFRs. Given the role of *Serpine1* in inflammation, tissue remodeling, and coagulopathy, this decreased accessibility may reflect impaired inflammatory reprogramming of MDSCs during sepsis.

A Class 7 example is the *Vcan* gene (**Figure 3G-I**) that encodes Versican, an extracellular matrix component that is upregulated during sepsis and thought so have some immune regulatory properties.^35,36^ Similar to Class 5 and 6 genes, the *Vcan* promoter in MDSCs from young or naïve female mice had no large NFRs, only displaying linker accessibility in arrays of randomly positioned nucleosomes (**Figure 3G**). However, as was observed for Class 5 and 6 loci, in MDSCs from older septic females, this characteristic linker accessibility diminished. Average chromatin accessibility across the *Vcan* promoter revealed a modest but consistent decrease in septic conditions, particularly in older, but not young, adult female mice (**Figure 3H**). Male adult septic mice (young and older) showed reduced average accessibility across the *Vcan* promoter, although the effect was less pronounced than that observed in older adult septic females. Conversely, young female and naïve adult male mice showed relatively higher accessibility, suggesting age- and sex-specific regulation. Again, no true DNA demethylation was observed at this locus and all subsequent loci; instead, only decreased accessibility at CCG sites was detected (**Figure 3I**).

#### 3. Promoters with constitutive NFRs that largely remain refractory to sepsis (Class 8)

Class 8 gene promoters showed high accessibility, with prominent NFRs encompassing the TSSs in all four conditions. However, these promoters illustrated an age-dependent, relatively modest loss of NFR accessibility in MDSCs from only older septic female mice, where overall NFR architecture and presumably transcriptional activity was largely preserved. Class 8 promoters generally exhibited low, baseline DNA methylation levels, reflective of their active status.

An example of Class 8 is the *Cd274* gene (**Figure 4A-C**), encoding the immune-checkpoint inhibitor PD-L1, which in MDSCs plays a direct role in suppressing T-cell responses.^38^ The *Cd274* promoter exhibited the highest levels of NFR accessibility in MDSCs from young septic mice and in naïve adult controls. MDSCs from older adult septic females showed a modest reduction in NFR accessibility but retained NFR structure, consistent with ongoing transcriptional activity. Average accessibility at the *Cd274* locus was modestly decreased in MDSCs from older adult septic female mice compared to naïve controls and young septic mice (**Figure 4B**). MDSCs from septic animals, particularly older individuals, showed a modest decrease in accessibility. Excluding DNA methylation of accessible CCG sites, only basal CpG methylation levels were detected within *Cd274* regulatory sequences, consistent with permissive chromatin and potential gene expression (**Figure 4C**).

**Figure 4.**
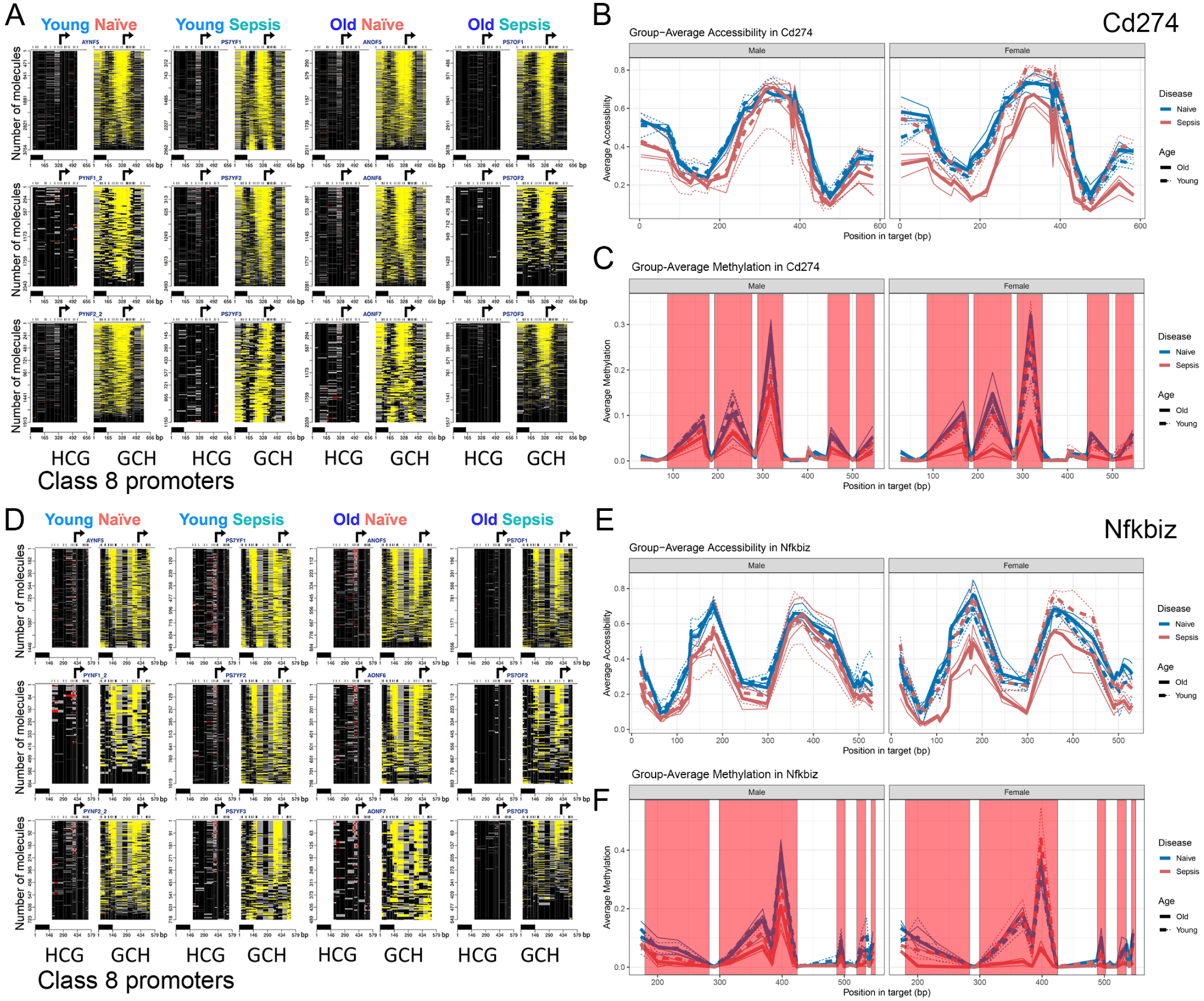
Class 8 promoters constitutively harbor well-defined NFRs across all conditions, which partially lose accessibility specifically in Female Older Sepsis (A) HCG and GCH plots for *Cd274* promoter show prominent NFRs across all conditions. The figure is annotated as in Figures 1 and 2. (B) Group-averaged chromatin accessibility (GpC methylation) across the same region, plotted separately for males (left) and females (right). (C) Group-averaged endogenous CpG methylation levels across the promoter. (D-F) Equivalent plots for *Nfkbiz* promoter.

Other Class 8 genes, such as *Nfkbiz* (encoding the NF-κB inhibitor IκBζ; **Figure 4D-F**), *Vdr* (vitamin D receptor), *Il4ra* (IL-4 receptor α chain, which has been shown to be important to MDSC-suppressive activity),^39^ and *Atf6* (ATF6, a transcription factor that activates target genes for the unfolded protein response) all followed a similar pattern with NFRs present in all conditions except in MDSCs from older septic female mice. In each case, MDSCs from the older female septic group showed baseline CpG methylation (omitting CCG) and a modest loss of promoter accessibility that is unlikely to impair transcriptional activity.

#### 4. Constitutively accessible promoters that maintain open chromatin in all conditions (Class 9)

Finally, a small subset of promoters (Class 9) remained highly accessible across all conditions, largely unaffected by sepsis, age, or sex. Class 9 promoters had pre-existing, prominent nucleosome-free regions in naïve MDSCs and retained these open states after the stress of sepsis. They also displayed low basal DNA methylation, consistent with active promoters. A clear example is the *Tet2* promoter, which encodes the DNA demethylation enzyme Tet2 (**Figure 5**). The *Tet2* promoter showed prominent NFRs in all samples across all conditions indicating constitutive chromatin accessibility. CpG methylation could not be assessed because out of 35 HGCs in the *Tet2* promoter amplicon all are CCGs.

**Figure 5.**
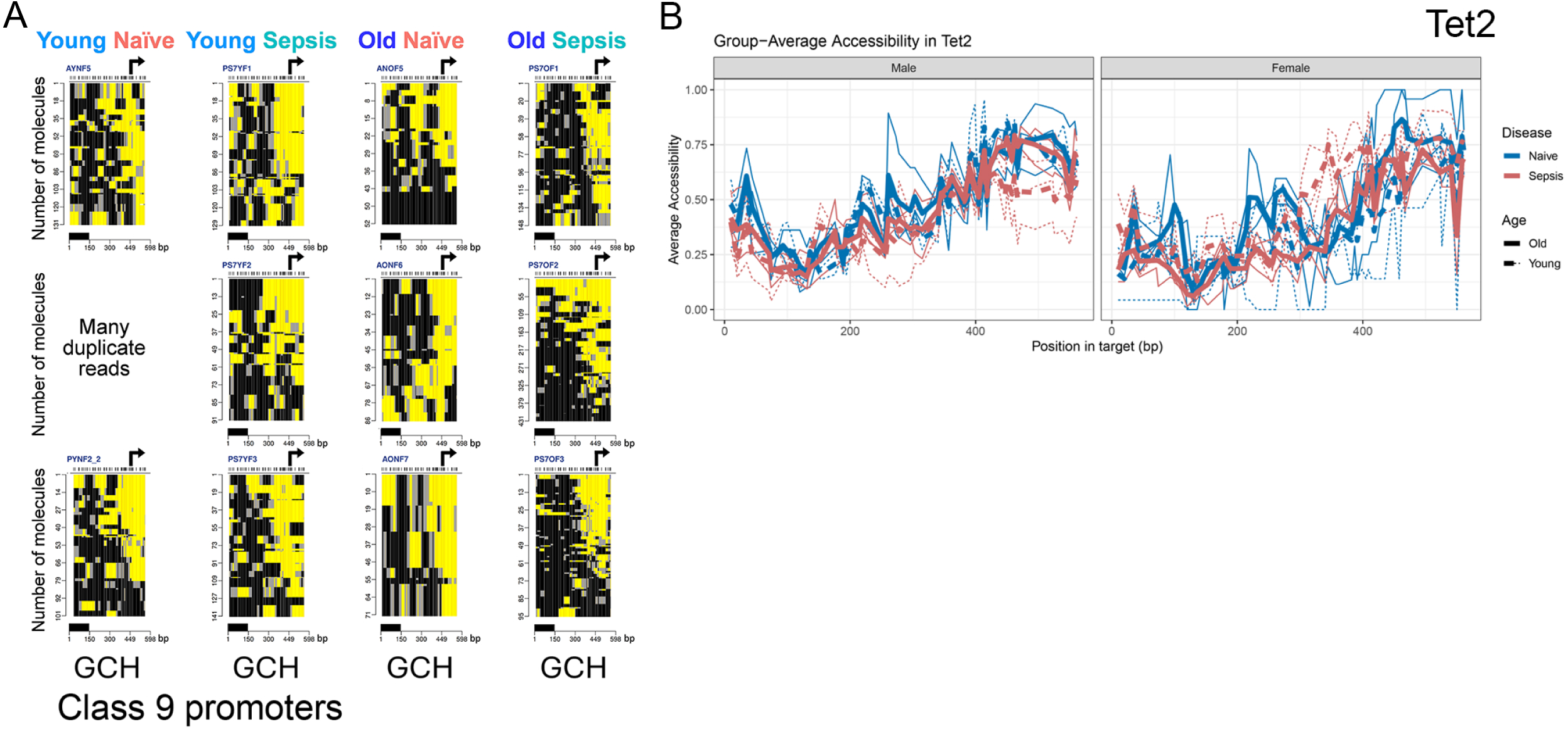
Class 9 promoters are constitutively open in sepsis (A) HCG and GCH plots for the *Tet2* promoter show strong, stable NFRs in all female MDSCs. The figure is annotated as in Figures 1 and 2. (B) Group-averaged chromatin accessibility (GpC methylation) across the same region, plotted separately for males (left) and females (right). No group-averaged endogenous CpG methylation in shown because all HCG sites are CCG.

### Single-cell transcriptomics support epigenetic promoter class distinctions in MDSC subsets

To determine whether promoter accessibility classes defined by MAPit-FENGC corresponded to functional gene expression patterns, we analyzed single-cell RNA-seq data from CD11b^+^Gr1⁺ splenic leukocytes isolated from young and older, male and female adult mice under naïve or septic conditions, collected in the Companion Paper (Rodhouse et al.). Data were stratified into three MDSC subtypes: early MDSCs (E-MDSC), monocytic MDSCs (M-MDSC), and granulocytic MDSCs (PMN-MDSC) based on gene expression patterns. The gene *S100a9*, assigned as a Class 1 promoter and characterized by widespread NFR formation in all septic groups, was induced across all three MDSC subsets in response to sepsis, regardless of age or sex (**Figure 6**). This transcriptional profile matches the broadly accessible chromatin configuration observed epigenetically. Similarly, the Class 3 promoter *S100a8*, which formed sepsis-induced NFRs in young and older mice, displayed upregulation in E-MDSCs and PMN-MDSCs during sepsis. In contrast, a Class 6 promoter *Ccl5*, which remained epigenetically inaccessible in all conditions, showed only modest and selective transcriptional induction in a subset of MDSCs, primarily E- and M-MDSCs, in young male septic mice (**Figure 7)**. These observations are consistent with a transcriptionally repressed or enhancer-driven regulatory program.

**Figure 6.**
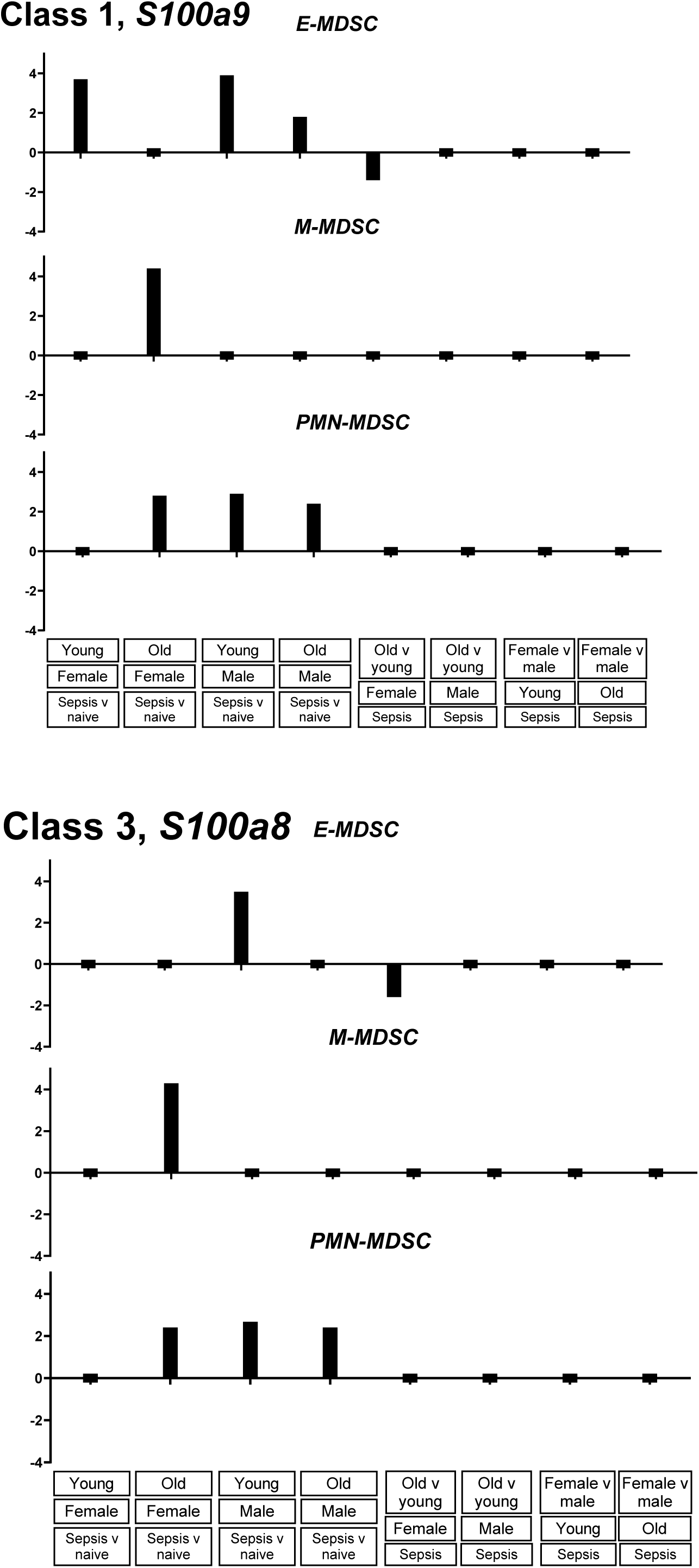
Class 1 and Class 3 genes show consistent sepsis-induced transcriptional activation across MDSC subsets Bar plots log_2 fo_ld change in expression for *S100a9* (Class 1) and *S100a8* (Class 3) across E-MDSC, M-MDSC, and PMN-MDSC populations (conditions and comparisons as labeled) derived from single-cell RNA-sequencing analysis of splenic Cd11b^+^ Gr1⁺ cells.

**Figure 7.**
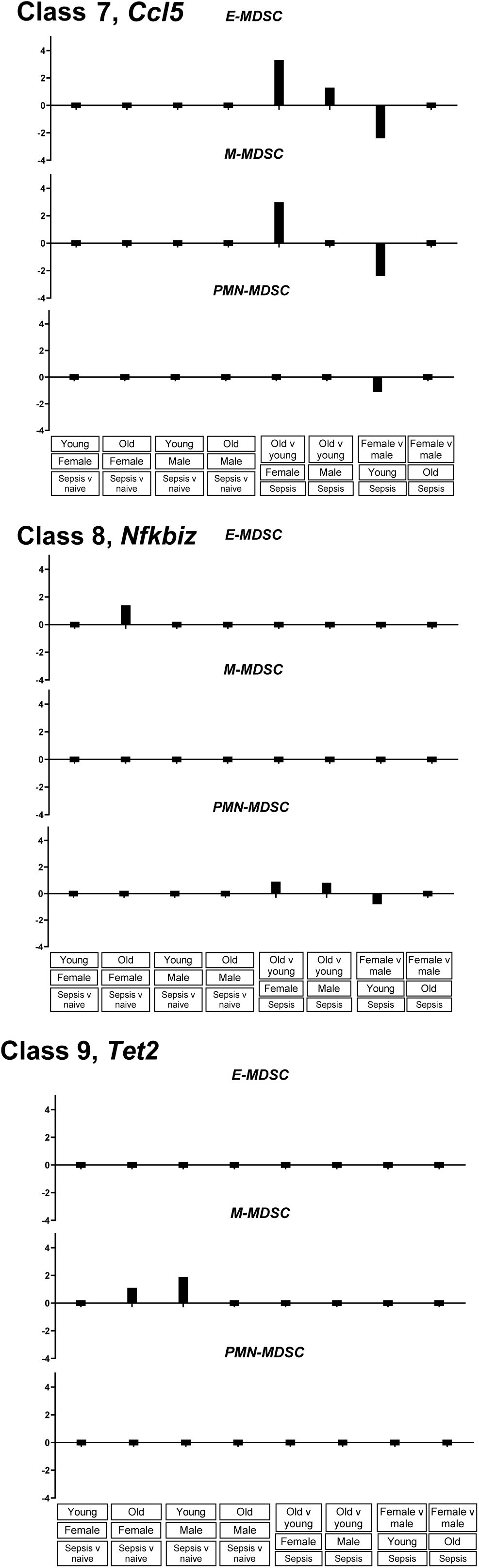
Expression of Class 7, Class 8 and Class 9 genes across MDSC subtypes reveals repression and age-dependent silencing Bar plots depict log_2 fo_ld change in gene expression (conditions and comparisons as labeled) for *Ccl5* (Class 7), *Nfkbiz* (Class 8) and *Tet2* (Class 9),derived from single-cell RNA-seq analysis of splenic Cd11b^+^ Gr1⁺ cells. Data are shown separately for early MDSCs (E-MDSC), monocytic MDSCs (M-MDSC), and polymorphonuclear MDSCs (PMN-MDSC).

In contrast, *Nfkbiz*, assigned to Class 8 based on selective modest promoter closure in “Aged Sepsis,” showed relatively low but detectable expression across MDSC subsets. While MAPit-FENGC revealed loss of promoter accessibility specifically in aged sepsis mice, scRNA-seq analysis did not show consistent transcriptional silencing in this group. Instead, modest expression changes were observed in PMN-MDSCs and E-MDSCs, particularly in young female septic mice. *Tet2*, a representative Class 9 gene whose promoter remained consistently accessible across all MAPit-FENGC conditions, exhibited modest transcriptional activation in scRNA-seq, primarily during sepsis in M-MDSCs. Despite the stable chromatin accessibility across all groups, expression was low or unchanged in E-MDSCs and PMN-MDSCs.

## DISCUSSION

Our study provides a high-resolution, single-molecule epigenetic map of splenic MDSCs in a murine model of surgical sepsis, revealing how host age and sex shape chromatin accessibility and DNA methylation in this immunoregulatory cell type. Using MAPit-FENGC, we identified nine distinct promoter classes across multiple immune-relevant loci, encompassing a spectrum of response classes ranging from sepsis-induced chromatin opening to age-specific silencing. These epigenetic patterns were tightly linked to host sex, age, and sepsis exposure and corresponded to changes in transcription, as assessed by scRNA-seq.

We discovered an age-dependent epigenetic repression of key immunoregulatory genes (e.g., *Cd274*, *Nfkbiz*, *Il4ra*) in older septic mice (Class 8). These promoters exhibited basal CpG methylation and robust promoter NFR accessibility in young or naïve mice. Older septic females showed a modest loss of accessibility but retention of NFR structure, consistent with ongoing transcriptional activity. Therefore, nucleosome repositioning emerges as the dominant mechanism of transcriptional repression in MDSCs from aged hosts. The selective, partial closure of immune checkpoint and anti-inflammatory loci (e.g., *Cd274*, *Il10*) may impair key MDSC functions (T-cell suppression and inflammation resolution), which rely on PD-L1 and IL-10 expression. In aged septic hosts, epigenetic repression of these genes may render MDSCs less effective or dysregulated, contributing to the persistent immune dysfunction and chronic critical illness seen in this population.^16,21^

In contrast, sepsis-induced chromatin opening was observed in promoter classes associated with canonical MDSC effector genes (Classes 1-4). Promoters such as *S100a9*, *S100a8*, *Nos2*, and *Ptgs2* gained accessibility in septic MDSCs, often with concurrent demethylation, consistent with transcriptional activation.^29,40^ The magnitude of these chromatin changes varied by sex and age. For example, Class 1 (*S100a9*) was accessible across all septic groups but showed particularly robust NFR formation in older females. These findings suggest that both age and sex can influence *S100a9* expression and chromatin accessibility, potentially contributing to the observed robust NFR formation at the *S100a9* promoter in older females during sepsis. Research has demonstrated that chromatin accessibility at the *S100a9* promoter is regulated during monocyte differentiation and is influenced by transcription factors such as C/EBPδ. In particular, C/EBPδ has been shown to bind within the *S100a9* promoter region, affecting its accessibility and expression levels.^41^ Age-related changes in immune function (“inflammaging”) have been associated with alterations in inflammatory markers, including S100a9. For example, a study investigating the impact of age and sex on inflammatory markers found that aging is associated with higher expression of *S100a9* and components of the NLRP3 inflammasome independent of sex^42^. In the context of systemic lupus erythematosus (SLE), S100a9 has been identified as a molecule integral in determining sex-specific immune responses,^30^ which may be relevant to understanding its regulation in other inflammatory conditions like sepsis. The *S100a8* and *S100a9* promoters (Class 1 and 3 in MAPit) exhibit universal sepsis-induced opening, particularly in aged females, and these genes are also among the top differentially expressed genes in our scRNA-seq dataset (in the accompanying manuscript, Rodhouse et al.). This gene pair is pivotal in MDSC activation, trafficking, and suppressive function. The robust accessibility and expression of *S100a8*/*a9* in both studies supports their role as central, sepsis-responsive effectors and potential biomarkers of MDSC reprogramming.

Class 2 promoters opened only in older females and males, but not young females highlighting a female-specific age threshold for epigenetic activation at specific loci. For example, CXCR2 is a chemokine receptor primarily expressed on neutrophils and plays a crucial role in mediating their recruitment to sites of infection, essential for survival in sepsis. Excessive or dysregulated CXCR2 signaling can contribute to tissue damage and organ failure,^43^ and the observed increased accessibility of the CXCR2 gene in older females and males may suggest an age-related epigenetic regulation leading to enhanced expression.

The identification of stably closed promoters (Class 5) and constitutively open loci (Class 9) may further describe the epigenetic plasticity in MDSCs. Class 5 genes (e.g., *Lyz1*, *F7*) remained silenced through hypermethylation and packaging into arrays of randomly positioned nucleosomes regardless of condition. The refractoriness of Class 5 promoters to opening indicates they are epigenetically “locked down” in splenic MDSCs. These genes remain transcriptionally inactive even in sepsis, suggesting they are either irrelevant to the MDSC sepsis response or otherwise tightly repressed in this cell type.

Classes 6 and 7 highlight additional promoters shut down in MDSCs. Although these promoters had less DNA methylation than Class 5 promoters, they nevertheless remained epigenetically repressed as gauged by inability to form large NFRs following CLP sepsis challenge. Like Class 5, the chromatin of Class 6 and 7 promoters consists of randomly positioned nucleosomes with accessible linker DNA. Across Class 6 and 7 promoters, we frequently observed that all MDSC samples from older septic mice lacked NFRs, whereas one or more of the other conditions retained an accessible configuration, suggesting age-specific chromatin condensation at these promoters.

Across Class 6 and 7 promoters, all MDSC samples from older septic mice consistently lacked NFRs characteristic of active promoters, while one or more of the other conditions retained localized linker accessibility. This pattern indicates age-specific chromatin condensation within promoter regions that are normally permissive, rather than a global loss of linker accessibility. As with all Class 5-7 promoters, these loci remain devoid of canonical NFRs, distinguishing them from the open promoter architectures observed in Classes 1-4.

Class 8 includes several immune-related genes (e.g., *Cd274*, *Il4ra*, *Nfkbiz*) whose partially diminished accessibility in older survivors of sepsis might contribute to impaired or altered immune regulation in this cohort. Notably, several Class 5–8 promoters, including *Cd274* (PD-L1), *Il10*, and *Nfkbiz*, exhibited significant chromatin closure selectively in MDSCs from older septic mice. This pattern suggests a dominant role for nucleosome occupancy in regulating gene expression in aged hosts. Supporting this, our companion transcriptomic study (Rodhouse et al.) demonstrated that these genes were only modestly or minimally induced at the RNA level in older MDSC populations following sepsis, particularly in M-MDSCs. These findings support the concept that age-associated epigenetic silencing of immunoregulatory loci may cause persistent immune dysfunction in older sepsis survivors, and that transcriptional non-responsiveness may be epigenetically preconditioned in this cohort.

Class 9 promoters (e.g., *Tet2*) were persistently accessible across all groups, suggesting these are “core” MDSC housekeeping genes that remain transcriptionally poised or active even in the face of sepsis-induced stress, are simply important for the development of this cell type (the demethylation enzyme TET2 is highly expressed in hematopoietic stem and progenitor cells and has been shown to be essential in regulating MDSC numbers in tumor models^44^ and models of emergency myelopoiesis post-LPS-challenge).^45^ Generally, Class 9 genes appear to be core functional genes whose promoter chromatin is stably open as part of the identity or necessary function of MDSCs. Their invariant accessibility contrasts the dynamic changes seen in Classes 1-8. The presence of Class 9 promoters underscores that some key promoters remain epigenetically unaltered and continuously active throughout sepsis. Maintenance of a large NFR at the *Tet2* promoter in all conditions exemplifies this stability. We did not observe any loss of accessibility at Class 9 loci, reinforcing that these promoters are resilient to both inflammatory stress and host factors.

These context-specific activation patterns suggest that MDSC-mediated immunosuppression may arise *via* different regulatory programs in males versus females, and in young versus older hosts. This observation aligns with our prior data on age- and sex-specific differences in MDSC transcriptomes and function,^17,19^ and may provide mechanistic insight into how such differences are epigenetically encoded.

Importantly, single-cell transcriptomics supported many of the epigenetic patterns described. Genes in Classes 1 and 3 showed increased expression in MDSC subsets concordant with sepsis-induced chromatin opening. In contrast, Class 8 genes (e.g., *Nfkbiz*, *Il4ra*) showed partial decoupling between accessibility and expression in older septic MDSC raising the possibility of additional regulatory checkpoints, such as enhancer silencing or post-transcriptional control, particularly in aged immune cells.^13,26^ These findings underscore the importance of integrating multiple levels of gene regulation (chromatin state, methylation, and transcription) when evaluating MDSC behavior in critical illness. Collectively, these scRNA-seq results support the biological relevance of the MAPit-defined promoter classes, illustrating how chromatin accessibility states correspond to transcriptional outcomes across sepsis, sex, and age in MDSC subsets.

The epigenetic heterogeneity observed here helps explain the divergent immune phenotypes/endotypes that we^46,47^ have defined in sepsis survivors, and provides a foundation for understanding MDSC dysfunction in aging. In particular, the age-specific silencing of checkpoint and anti-inflammatory loci may promote persistent immune suppression, a hallmark of chronic critical illness and poor sepsis recovery.^5,16^ Conversely, sex-biased promoter accessibility could underlie known disparities in sepsis outcomes between males and females.^18^ These mechanistic insights may guide future strategies to therapeutically reprogram MDSCs using epigenetic modulators such as histone deacetylase (HDAC) inhibitors, bromodomain inhibitors, or DNA methyltransferase inhibitors, many of which are already being explored in oncologic and inflammatory contexts.^7,26^

Moreover, promoter classes defined here could serve as biomarkers of MDSC-state or targets for selective modulation. For instance, genes in Classes 1-3 may be used to identify actively suppressive MDSCs post-sepsis, while Class 8 promoters might be harnessed to identify aged immune populations with impaired regulatory function. The stability of Class 9 loci suggests they may serve as internal controls or indicators of MDSC identity in future epigenetic studies.

Our findings provide mechanistic insight into how sepsis may drive divergent epigenetic fates in MDSCs that reflect hallmarks of both trained immunity and tolerance. The persistent chromatin accessibility observed at Class 1-3 promoters (e.g., *S100a9*, *Nos2*, *S100a8*), particularly in older or female hosts, suggests a form of innate immune training, where prior inflammatory exposure leaves a molecular imprint that primes these loci for rapid reactivation. This is consistent with studies showing that chromatin remodeling at promoters of inflammatory genes underlies the trained phenotype in monocytes and macrophages (reviewed in^48^). Conversely, the selective silencing of promoters in Classes 6-8, including key regulatory genes such as *Cd274* and *Il10*, supports the emergence of an epigenetic tolerance program, particularly in aged MDSCs. These loci become refractory to activation despite hypomethylation, implicating nucleosome repositioning and chromatin compaction as dominant suppressive mechanisms. The coexistence of both accessibility gains and repressive remodeling across promoter classes indicates that sepsis induces a divergent epigenetic landscape in MDSCs, poising some genes for increased responsiveness while silencing others. These dual trajectories may underlie the paradoxical state of immune hyperinflammation coupled with functional suppression seen in sepsis survivors and reflect an imprint of the inflammatory milieu on the myeloid epigenome that varies by age and sex.

### Limitations

This study focused on a pre-selected set of immune and MDSC-relevant promoters, limiting discovery of unanticipated regulatory regions or enhancer-driven transcription. Additionally, while we profiled Gr1⁺CD11b⁺ splenocytes, these include multiple MDSC subtypes and potentially other myeloid cells. Although scRNA-seq helped disaggregate subtype-specific effects, future studies using single-cell epigenomics (e.g., scATAC-seq or scCUT&Tag) could more finely resolve MDSC heterogeneity.^49^ Finally, the MAPit-FENGC dataset provides high-resolution data at selected promoters—the approach can be readily extended to capture distal regulatory elements, which may play significant roles in aging-related immune dysfunction.

## Resource Availability

Requests for further information and resources should be directed to and will be fulfilled by the lead contact, Michael Kladde, PhD. This study did not generate new unique reagents. Data pending submission to GEO.

## Acknowledgments

This work was supported, in part, by the following grants:

National Institutes of Health T32GM008721, T32HL160491, RF1NS128626, R21AG087039 Department of Defense, Defense Threat Reduction Agency Grant 11912108

## Author Contributions

Conceptualization: MPK, PAE

Formal analysis: RLB, DBD, MPLG, JOB, MZ, FX, GC, RM, MPK

Funding acquisition: PAE, MPK, GC, PC

Investigation: MPLG, JOB, MPK

Methodology: MPK, MZ

Project administration: MPK

Supervision: PAE, LLM, RM, MPK

Visualization: RM, MPK

Writing – original draft: AMC, DBD, CER, PAE, RM, MPK

Writing – review & editing: CER, AMC, MH, MPG, JOB, RLB, LB, AMM, FX, GC, SMW, CEM, LLM, PAE, RM, MPK

## Methods

### Experimental Model

#### Animals

Young and older adult female (50%) and male (50%) mice underwent our intra-abdominal sepsis model of cecal ligation and puncture (CLP) followed by daily chronic stress (DCS). Approximately eight to 12 mice were used per experimental group. Mice were euthanized seven days post CLP + DCS. Experimental mice were euthanized in compliance with predetermined IACUC Body Condition Score (BCS) criteria. A modification to this included the following: greater than or equal to 20% weight loss from baseline or from age-matched controls if the animals are still growing during the study or greater than or equal to 20% weight loss from baseline in older mice who have a BCS of greater than 2 in the first five days after CLP, or greater than 40% weight loss from baseline in older mice with a BCS greater than 2 after five days following CLP. Based on these criteria, no mice had to be euthanized for excessive weight loss during the course of the study.

#### Intra-Abdominal Sepsis Model

CLP was conducted under isoflurane anesthesia we have previously described^50^. The cecum was ligated 1 cm from its tip and a 22- or 25-gauge needle was used to puncture the cecum depending on the desired experimental mortality. Buprenorphine analgesia was provided for 48 hours post surgery. Antibiotics (imipenem monohydrate; 25 mg/kg diluted in 1 mL 0.9% sodium chloride [NaCl]) were administered two hours post-CLP then continued for 72 hours. DCS was conducted as previously described^51^. Briefly, this involved placing mice in weighted plexiglass animal restraint holders (Kent Scientific, Torrington, CT) for two hours daily starting the day after CLP. The purpose was to simulate the ICU environment, where patients are often bedbound with limited mobility. MDSC were isolated using a CD11b^+^Gr1^+^ isolation (EasySep™ Mouse MDSC (CD11b+Gr1+) Isolation Kit, StemCell, Cambridge, MA, USA) according to manufacturer’s instructions and as we have described previously^21^.

### Methods Details

#### Single-cell RNA sequencing (scRNA-seq) analysis

Splenocytes were obtained and counts were quantified using a Cellometer™ Auto 2000 Cell Viability Counter (Nexcelom Bioscience, Lawrence, MA). One million cells with a minimum of 85% cell viability were obtained for scRNAseq using 10X Genomics chemistry (Chromium X instrument, Pleasonton, CA). as described in the companion paper (Rodhouse *et al*).

#### Methyltransferase Accessibility Protocol for Individual Templates combined with Flap-Enabled Next-Generation Capture (MAPit-FENGC) epigenetic analysis

MAPit-FENGC is a validated method that allows for simultaneous single-molecular level analysis of methylation and accessibility at target promoter and enhancer regions^27^. One million cells from each mouse were washed with ice-cold PBS, pH 7.2. Cells were centrifuged at 1,000 *x g* for five min then washed in ice-cold cell resuspension buffer (20 mM HEPES pH 7.5, 70 mM NaCl, 0.25 mM EDTA pH 8.0, 0.5 mM EGTA pH 8.0, 0.5% glycerol, with freshly supplemented 10 mM DTT and 0.25 mM PMSF). Cells were centrifuged at 1,000 *x g* for five min before being resuspended with 92 μL of resuspension buffer with 0.5 % digitonin. Cells were then stained with trypan blue to ascertain 100 % permeabilization. The cells were then treated with 100 U M.CviPI GpC methyltransferase (100 U / million cells; New England Biolabs, M0227B-HI) with fresh 160 160 µM S-adenosyl-L-methionine for 15 min at 37°C. The reaction was terminated using an equal volume of 10 mM EDTA, 100 mM NaCl, and 1% (w/v) SDS followed by a quick vortex at medium speed. The nuclei were treated with 100 µg/ml RNase A for 30 min at 37°C followed with 100 µg/ml proteinase K treatment at 50°C overnight. Extraction of genomic DNA was conducted using phenol-chloroform-isoamyl alcohol (25:24:1, v/v) phase separation, which was followed by ethanol precipitation, then resuspension in molecular-grade H_2O_. The full FENGC methodology can be found at Zhou et al. (2022)^27^. Target gene promoters and enhancers were located using the transcription start site (TSS) region and encode annotations from the mm10 genome assembly. Flap oligos 1 and 2 as well as the nested oligos 3 are catalogued in **Supplementary Material 1**. All three oligos (4 nmole each) were ordered as 0.3-mL each as salt-free oligos in 96 well format (Eurofins Genomics, KY, USA). Purified amplicons were submitted to the University of Florida Interdisciplinary Center for Biotechnology Research (UF-ICBR, RRID:SCR_019152) for SMRT bell library construction. Sequencing was run on PacBio SEQUEL IIe (City, State) instrument by UF-ICBR. The library pool was loaded at 120 pM, using diffusion loading and 20- to 30-h movies with HiFi generation and demultiplexing. Sequencing Kit 2.0 (PacBio, 101-389-001) and Instrument Chemistry Bundle Version 11.0 were used with all other steps being performed using recommended protocol by the PacBio sequencing calculator. For epigenetic analysis, high-fidelity circular consensus sequencing (CCS) was generated using default parameters except for a setting of ≥ 5 single polymerase read passes. CCS reads were aligned to the reference sequences using the python reAminator pipeline^52^. Cut-offs to 95% conversion rate and 95% length of reference sequences alignments were applied. To distinguish endogenous CG methylation and M.CviPI-probed GC methylation, the unambiguously GCG sites were removed from calculation of HCG and GCH methylation (where H is A, C, or T). Averaged heatmaps were constructed based on averaged percent HCG/GCH per nucleotide post-methylscaper analysis^53^. A total of 180 primers for selected promoter and enhancer regions of immune and trauma-related genes (**Supplemental Material 1**) were used to concurrently profile chromatin accessibility and methylation. To assess global epigenetic differences in CD11b^+^ Gr1⁺ MDSCs isolated from murine spleens, we used three types of data matrices: (1) endogenous CpG methylation (HCG sites), (2) GpC methylation representing chromatin accessibility (GCH sites), and (3) combined HCG + GCH profiles. For each sample, a matrix of percent methylation across covered sites was generated and filtered to retain loci with ≥25 informative reads. Principal component analysis (PCA) was conducted using base R’s prcomp() function without imputation. After quality filtering, each promoter target was filtered for fewer than 25 reads. Targets were not used in the PCAs if < 2 samples had data passing the filter.

---

**Table 1.** Gene Promoters Grouped by Epigenetic Class.

| Epigenetic Class | Gene Promoters |
| --- | --- |
| 1 | *S100a9* |
| 2 | *Ccl3, Cxcr2, Il1rl2, Lcn2, Lgals9, Nos2, Ptgs2, Rnase2a* |
| 3 | *Mmp8, Ros1, S100a8* |
| 4 | *Fyb* |
| 5 | *Dab2, Emp1, F7, Lyz1, Mmp19, Pmp22, Retn, Serpina1a, Vnn1, Vsig4* |
| 6 | *Ambp, Ccl5, Cd9, Cxcl3, Fabp7, Fgr, Gpnmb, Kdm6b, Lgals2, Mmp9, Saa3, Serpine1, Sparc* |
| 7 | *Car4, Ccl17, Gprc5b, Htr2a, Igkv12-44, Igkv12-46, Igkv4-69 , Il10, Mt2, Plac8, S100a10, Stab1, Tmem176a, Tmem176a, Vcan* |
| 8 | *Atf6, Cd274, Hdac3, Il4ra, Nfkbiz, Vdr* |
| 9 | *Jun, Tet2, Stat3* |

**Supplemental Fig 1:**
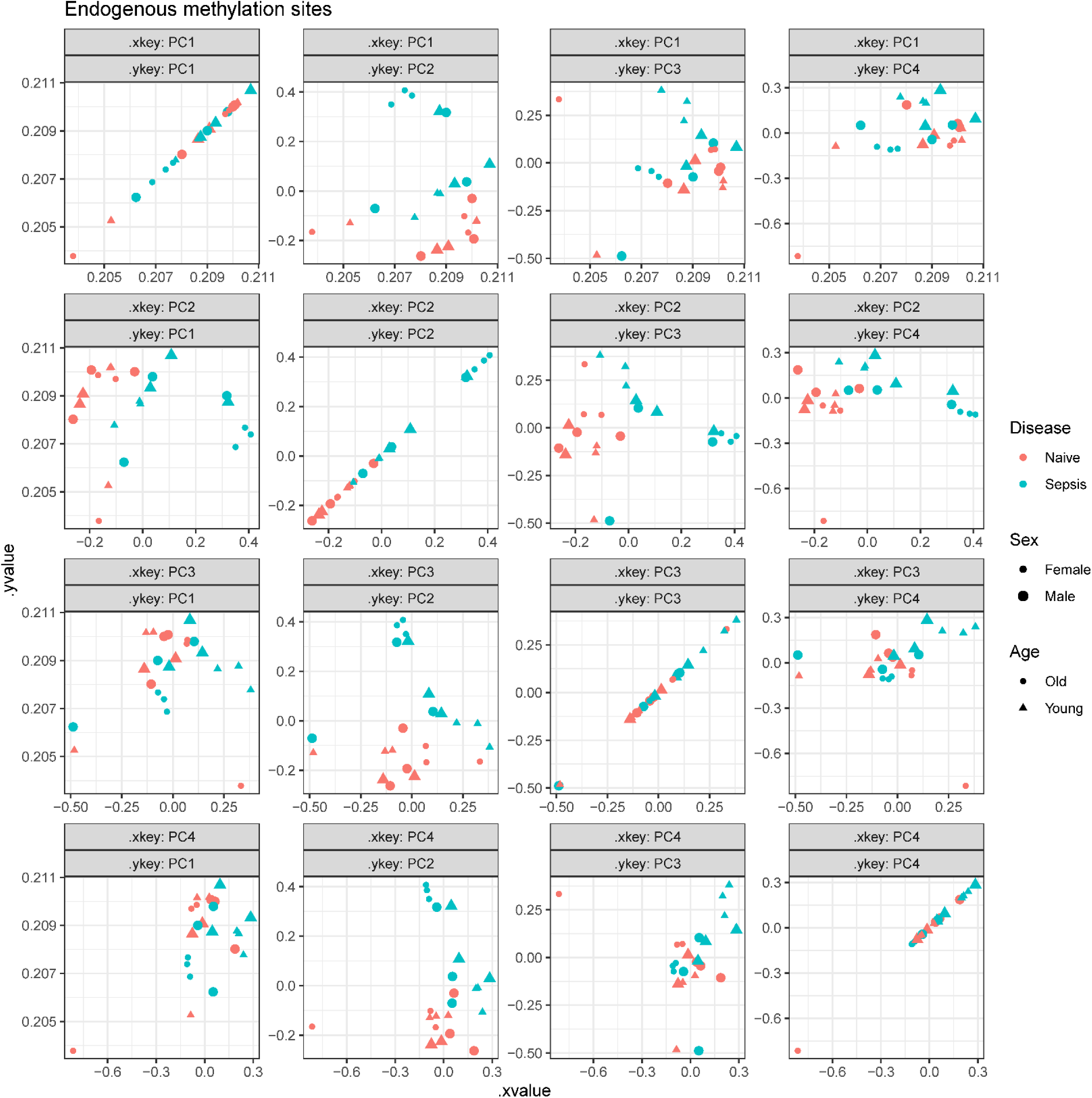
Principal component analysis (PCA) of endogenous CpG methylation (HCG) in splenic MDSCs. PCA was performed on the percent methylation of CpG (HCG) sites across all covered promoters in CD11b+ Gr1⁺ MDSCs from male and female mice, young and old, with or without sepsis. Each point represents a single mouse. Color indicates disease condition (blue = naïve, red = sepsis), shape denotes sex (circle = female, triangle = male), and shading differentiates age (light = young, dark = old). Principal components 1-4 are plotted in pairwise combinations to visualize clustering. Old septic females exhibit distinct separation, indicating strong methylome remodeling in response to sepsis.

**Supplemental Fig 2:**
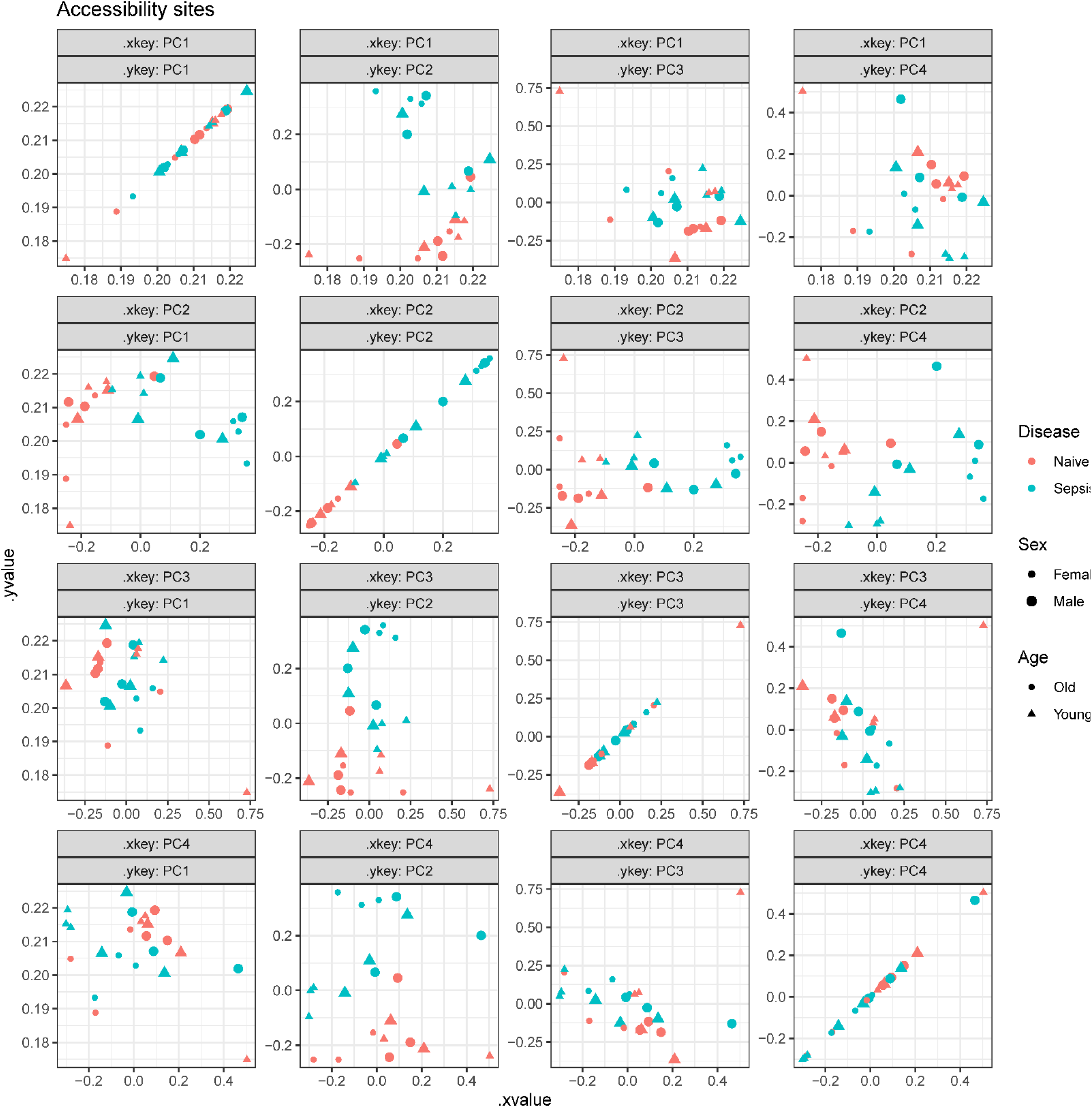
Principal component analysis (PCA) of GpC methylation (chromatin accessibility) in splenic MDSCs. PCA was performed using GCH methylation values from MAPit-FENGC to reflect promoter accessibility across MDSC samples. Each point represents a single mouse. Color indicates disease condition (blue = naïve, red = sepsis), shape denotes sex (circle = female, triangle = male), and shading differentiates age (light = young, dark = old). Separation by sepsis status is evident across multiple PCs, with the strongest divergence observed in female mice. These results suggest widespread chromatin remodeling following sepsis, particularly in females.

**Supplemental Table 1.**
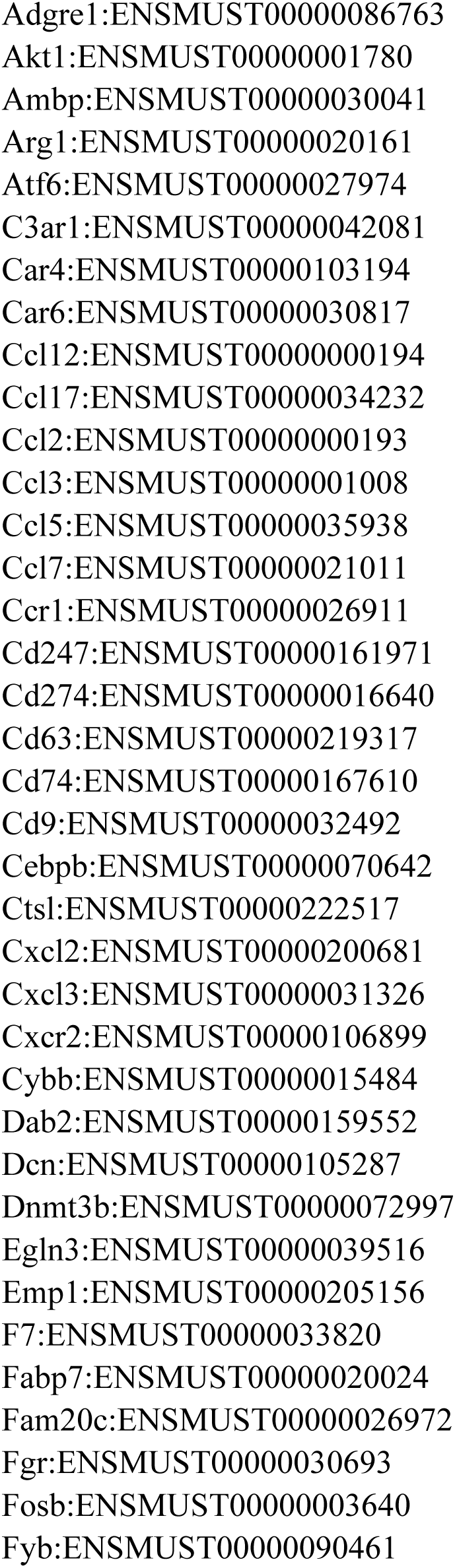
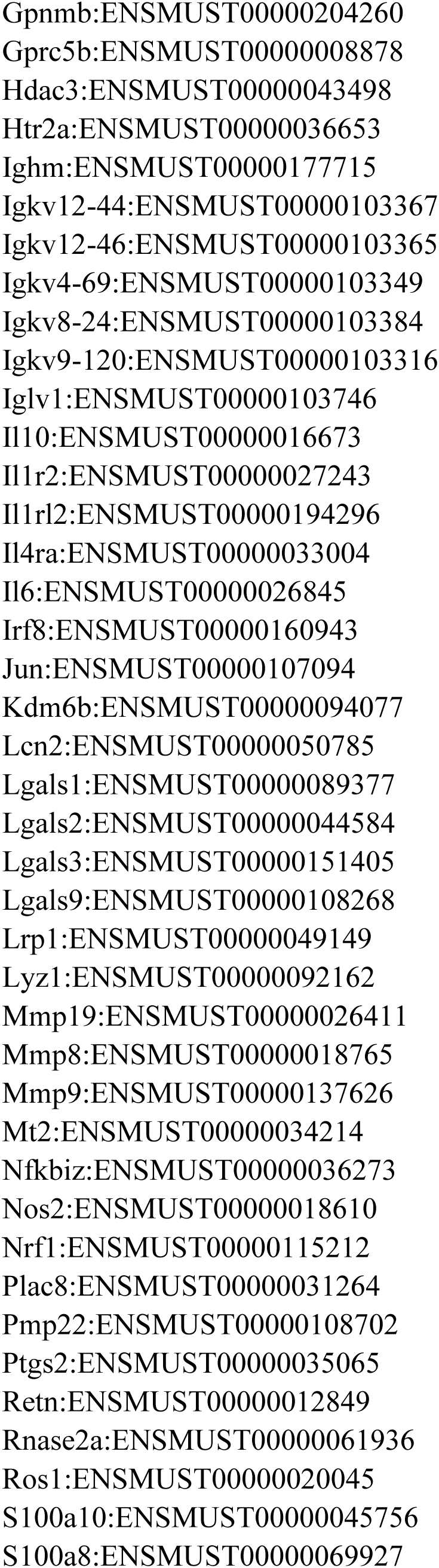
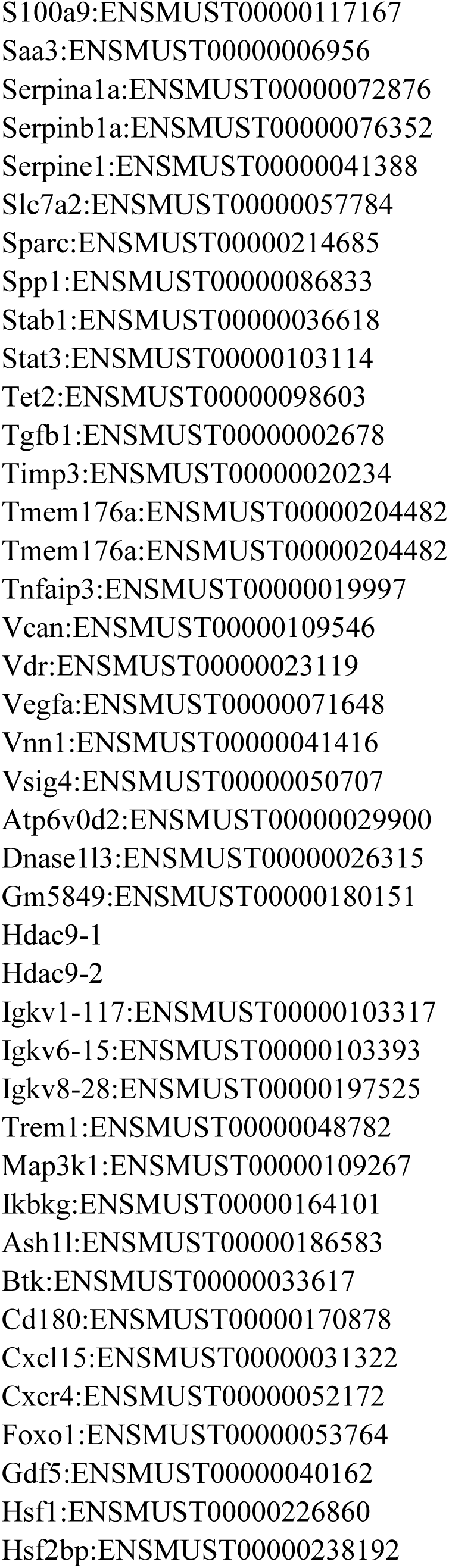
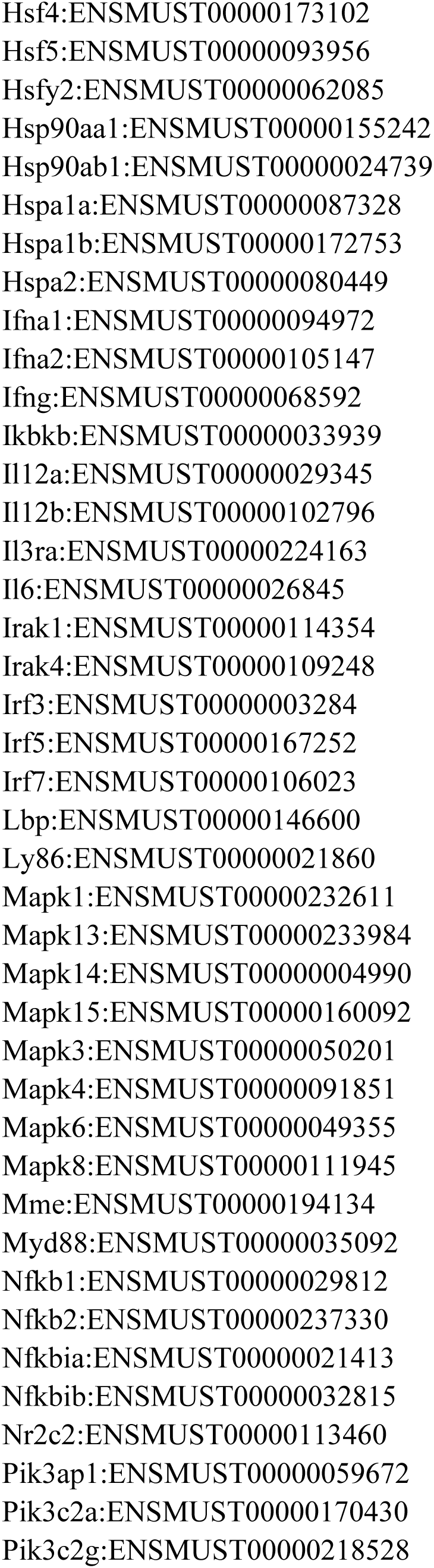
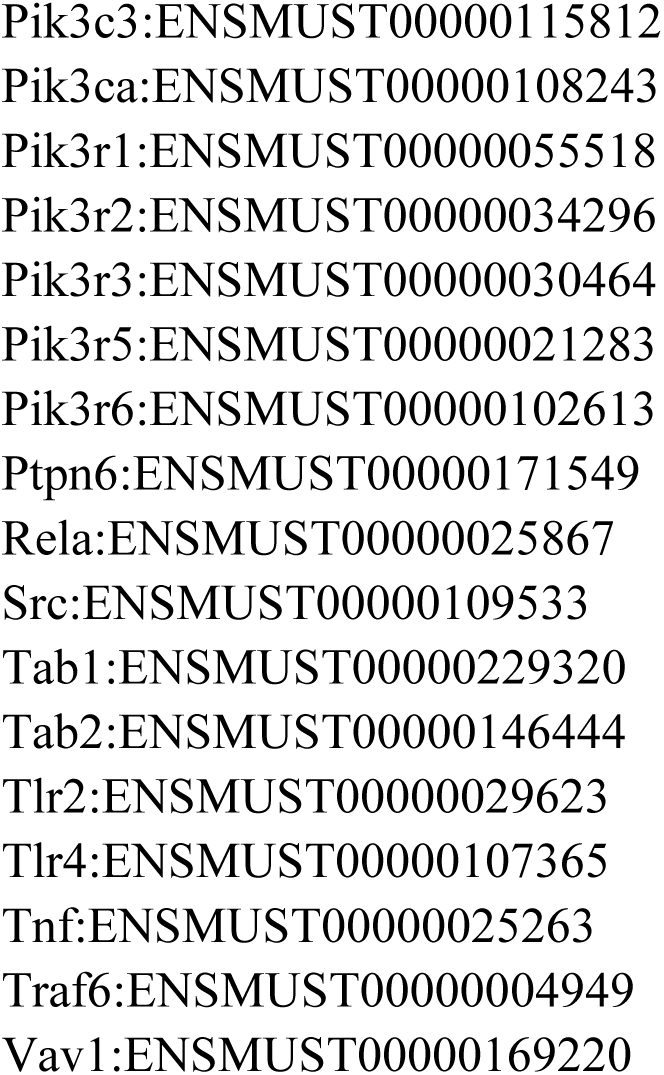
Gene and Primer List used for MAPit-FENGC.

## REFERENCES

1. Singer, M., Deutschman, C.S., Seymour, C.W., Shankar-Hari, M., Annane, D., Bauer, M., Bellomo, R., Bernard, G.R., Chiche, J.D., Coopersmith, C.M., et al. (2016). The Third International Consensus Definitions for Sepsis and Septic Shock (Sepsis-3). Jama 315, 801–810. 10.1001/jama.2016.0287.

2. Rhee, C., Dantes, R., Epstein, L., Murphy, D.J., Seymour, C.W., Iwashyna, T.J., Kadri, S.S., Angus, D.C., Danner, R.L., Fiore, A.E., et al. (2017). Incidence and Trends of Sepsis in US Hospitals Using Clinical vs Claims Data, 2009-2014. Jama 318, 1241–1249. 10.1001/jama.2017.13836.

3. Hollenbeak, C.S., Henning, D.J., Geeting, G.K., Ledeboer, N.A., Faruqi, I.A., Pierce, C.G., Thomas, C.B., and O’Neal, H.R., Jr. (2023). Costs and Consequences of a Novel Emergency Department Sepsis Diagnostic Test: The IntelliSep Index. Crit Care Explor 5, e0942. 10.1097/CCE.0000000000000942.

4. Herran-Monge, R., Muriel-Bombin, A., Garcia-Garcia, M.M., Merino-Garcia, P.A., Martinez-Barrios, M., Andaluz, D., Ballesteros, J.C., Dominguez-Berrot, A.M., Moradillo-Gonzalez, S., Macias, S., et al. (2019). Epidemiology and Changes in Mortality of Sepsis After the Implementation of Surviving Sepsis Campaign Guidelines. J Intensive Care Med 34, 740–750. 10.1177/0885066617711882.

5. Darden, D.B., Brakenridge, S.C., Efron, P.A., Ghita, G.L., Fenner, B.P., Kelly, L.S., Mohr, A.M., Moldawer, L.L., and Moore, F.A. (2021). Biomarker Evidence of the Persistent Inflammation, Immunosuppression and Catabolism Syndrome (PICS) in Chronic Critical Illness (CCI) After Surgical Sepsis. Ann Surg 274, 664–673. 10.1097/SLA.0000000000005067.

6. Brakenridge, S.C., Efron, P.A., Cox, M.C., Stortz, J.A., Hawkins, R.B., Ghita, G., Gardner, A., Mohr, A.M., Anton, S.D., Moldawer, L.L., and Moore, F.A. (2019). Current Epidemiology of Surgical Sepsis: Discordance Between Inpatient Mortality and 1-year Outcomes. Ann Surg 270, 502–510. 10.1097/sla.0000000000003458.

7. Polcz, V.E., Barrios, E.L., Larson, S.D., Efron, P.A., and Rincon, J.C. (2024). Charting the course for improved outcomes in chronic critical illness: therapeutic strategies for persistent inflammation, immunosuppression, and catabolism syndrome (PICS). Br J Anaesth 133, 260–263. 10.1016/j.bja.2024.05.005.

8. Gabrilovich, D.I., Bronte, V., Chen, S.H., Colombo, M.P., Ochoa, A., Ostrand-Rosenberg, S., and Schreiber, H. (2007). The terminology issue for myeloid-derived suppressor cells. Cancer Res 67, 425; author reply 426. 10.1158/0008-5472.CAN-06-3037.

9. Veglia, F., Sanseviero, E., and Gabrilovich, D.I. (2021). Myeloid-derived suppressor cells in the era of increasing myeloid cell diversity. Nat Rev Immunol 21, 485–498. 10.1038/s41577-020-00490-y.

10. Mira, J.C., Gentile, L.F., Mathias, B.J., Efron, P.A., Brakenridge, S.C., Mohr, A.M., Moore, F.A., and Moldawer, L.L. (2017). Sepsis Pathophysiology, Chronic Critical Illness, and Persistent Inflammation-Immunosuppression and Catabolism Syndrome. Crit Care Med 45, 253–262. 10.1097/CCM.0000000000002074.

11. Uhel, F., Azzaoui, I., Gregoire, M., Pangault, C., Dulong, J., Tadie, J.M., Gacouin, A., Camus, C., Cynober, L., Fest, T., et al. (2017). Early Expansion of Circulating Granulocytic Myeloid-derived Suppressor Cells Predicts Development of Nosocomial Infections in Patients with Sepsis. Am J Respir Crit Care Med 196, 315–327. 10.1164/rccm.201606-1143OC.

12. Mathias, B., Delmas, A.L., Ozrazgat-Baslanti, T., Vanzant, E.L., Szpila, B.E., Mohr, A.M., Moore, F.A., Brakenridge, S.C., Brumback, B.A., Moldawer, L.L., et al. (2017). Human Myeloid-derived Suppressor Cells are Associated With Chronic Immune Suppression After Severe Sepsis/Septic Shock. Ann Surg 265, 827–834. 10.1097/SLA.0000000000001783.

13. Barrios, E.L., Leary, J.R., Darden, D.B., Rincon, J.C., Willis, M., Polcz, V.E., Gillies, G.S., Munley, J.A., Dirain, M.L., Ungaro, R., et al. (2024). The post-septic peripheral myeloid compartment reveals unexpected diversity in myeloid-derived suppressor cells. Front Immunol 15, 1355405. 10.3389/fimmu.2024.1355405.

14. Darden, D.B., Bacher, R., Brusko, M.A., Knight, P., Hawkins, R.B., Cox, M.C., Dirain, M.L., Ungaro, R., Nacionales, D.C., Rincon, J.C., et al. (2021). Single-Cell RNA-seq of Human Myeloid-Derived Suppressor Cells in Late Sepsis Reveals Multiple Subsets With Unique Transcriptional Responses: A Pilot Study. Shock 55, 587–595. 10.1097/SHK.0000000000001671.

15. Hollen, M.K., Stortz, J.A., Darden, D., Dirain, M.L., Nacionales, D.C., Hawkins, R.B., Cox, M.C., Lopez, M.C., Rincon, J.C., Ungaro, R., et al. (2019). Myeloid-derived suppressor cell function and epigenetic expression evolves over time after surgical sepsis. Crit Care 23, 355. 10.1186/s13054-019-2628-x.

16. Efron, P.A., Brakenridge, S.C., Mohr, A.M., Barrios, E.L., Polcz, V.E., Anton, S., Ozrazgat-Baslanti, T., Bihorac, A., Guirgis, F., Loftus, T.J., et al. (2023). The persistent inflammation, immunosuppression, and catabolism syndrome 10 years later. J Trauma Acute Care Surg 95, 790–799. 10.1097/TA.0000000000004087.

17. Efron, P.A., Darden, D.B., Li, E.C., Munley, J., Kelly, L., Fenner, B., Nacionales, D.C., Ungaro, R.F., Dirain, M.L., Rincon, J., et al. (2022). Sex differences associate with late microbiome alterations after murine surgical sepsis. J Trauma Acute Care Surg 93, 137–146. 10.1097/ta.0000000000003599.

18. Rani, A., Barter, J., Kumar, A., Stortz, J.A., Hollen, M., Nacionales, D., Moldawer, L.L., Efron, P.A., and Foster, T.C. (2022). Influence of age and sex on microRNA response and recovery in the hippocampus following sepsis. Aging (Albany NY) 14, 728–746. 10.18632/aging.203868.

19. Mankowski, R.T., Thomas, R.M., Darden, D.B., Gharaibeh, R.Z., Hawkins, R.B., Cox, M.C., Apple, C., Nacionales, D.C., Ungaro, R.F., Dirain, M.L., et al. (2021). Septic Stability? Gut Microbiota in Young Adult Mice Maintains Overall Stability After Sepsis Compared to Old Adult Mice. Shock 55, 519–525. 10.1097/shk.0000000000001648.

20. Mankowski, R.T., Anton, S.D., Ghita, G.L., Brumback, B., Cox, M.C., Mohr, A.M., Leeuwenburgh, C., Moldawer, L.L., Efron, P.A., Brakenridge, S.C., and Moore, F.A. (2020). Older Sepsis Survivors Suffer Persistent Disability Burden and Poor Long-Term Survival. J Am Geriatr Soc 68, 1962–1969. 10.1111/jgs.16435.

21. Stortz, J.A., Hollen, M.K., Nacionales, D.C., Horiguchi, H., Ungaro, R., Dirain, M.L., Wang, Z., Wu, Q., Wu, K.K., Kumar, A., et al. (2019). Old Mice Demonstrate Organ Dysfunction as well as Prolonged Inflammation, Immunosuppression, and Weight Loss in a Modified Surgical Sepsis Model. Crit Care Med 47, e919–e929. 10.1097/ccm.0000000000003926.

22. Barter, J., Kumar, A., Stortz, J.A., Hollen, M., Nacionales, D., Efron, P.A., Moldawer, L.L., and Foster, T.C. (2019). Age and Sex Influence the Hippocampal Response and Recovery Following Sepsis. Mol Neurobiol 56, 8557–8572. 10.1007/s12035-019-01681-y.

23. Brakenridge, S.C., Efron, P.A., Stortz, J.A., Ozrazgat-Baslanti, T., Ghita, G., Wang, Z., Bihorac, A., Mohr, A.M., Brumback, B.A., Moldawer, L.L., and Moore, F.A. (2018). The impact of age on the innate immune response and outcomes after severe sepsis/septic shock in trauma and surgical intensive care unit patients. J Trauma Acute Care Surg 85, 247–255. 10.1097/TA.0000000000001921.

24. Raymond, S.L., Lopez, M.C., Baker, H.V., Larson, S.D., Efron, P.A., Sweeney, T.E., Khatri, P., Moldawer, L.L., and Wynn, J.L. (2017). Unique transcriptomic response to sepsis is observed among patients of different age groups. PLoS One 12, e0184159. 10.1371/journal.pone.0184159.

25. Nacionales, D.C., Szpila, B., Ungaro, R., Lopez, M.C., Zhang, J., Gentile, L.F., Cuenca, A.L., Vanzant, E., Mathias, B., Jyot, J., et al. (2015). A Detailed Characterization of the Dysfunctional Immunity and Abnormal Myelopoiesis Induced by Severe Shock and Trauma in the Aged. J Immunol 195, 2396–2407. 10.4049/jimmunol.1500984.

26. Gjerstorff, M.F. (2023). Epigenetic targeting of myeloid-derived suppressor cells: time to move into infectious diseases? Front Immunol 14, 1247715. 10.3389/fimmu.2023.1247715.

27. Zhou, M., Nabilsi, N.H., Wang, A., Gauthier, M.-P.L., Murray, K.O., Azari, H., Owens, W.S., Newman, J.R.B., Pardo-Palacios, F.J., Conesa, A., et al. (2022–11–09). Flap-enabled next-generation capture (FENGC): precision targeted single-molecule profiling of epigenetic heterogeneity, chromatin dynamics, and genetic variation. bioRxiv. 10.1101/2022.11.08.515732.

28. Knight, P., Gauthier, M.L., Pardo, C.E., Darst, R.P., Kapadia, K., Browder, H., Morton, E., Riva, A., Kladde, M.P., and Bacher, R. (2021). Methylscaper: an R/Shiny app for joint visualization of DNA methylation and nucleosome occupancy in single-molecule and single-cell data. Bioinformatics 37, 4857–4859. 10.1093/bioinformatics/btab438.

29. von Wulffen, M., Luehrmann, V., Robeck, S., Russo, A., Fischer-Riepe, L., van den Bosch, M., van Lent, P., Loser, K., Gabrilovich, D.I., Hermann, S., et al. (2023). S100A8/A9-alarmin promotes local myeloid-derived suppressor cell activation restricting severe autoimmune arthritis. Cell Rep 42, 113006. 10.1016/j.celrep.2023.113006.

30. Davison, L.M., Alberto, A.A., Dand, H.A., Keller, E.J., Patt, M., Khan, A., Dvorina, N., White, A., Sakurai, N., Liegl, L.N., et al. (2021). S100a9 Protects Male Lupus-Prone NZBWF1 Mice From Disease Development. Front Immunol 12, 681503. 10.3389/fimmu.2021.681503.

31. Bullock, K., and Richmond, A. (2021). Suppressing MDSC Recruitment to the Tumor Microenvironment by Antagonizing CXCR2 to Enhance the Efficacy of Immunotherapy. Cancers (Basel) 13. 10.3390/cancers13246293.

32. Santibanez, J.F. (2025). Myeloid-Derived Suppressor Cells: Implications in Cancer Immunology and Immunotherapy. Front Biosci (Landmark Ed) 30, 25203. 10.31083/FBL25203.

33. Wang, S., Zhao, X., Wu, S., Cui, D., and Xu, Z. (2023). Myeloid-derived suppressor cells: key immunosuppressive regulators and therapeutic targets in hematological malignancies. Biomark Res 11, 34. 10.1186/s40364-023-00475-8.

34. Ibrahim, A.A., Fujimura, T., Uno, T., Terada, T., Hirano, K.I., Hosokawa, H., Ohta, A., Miyata, T., Ando, K., and Yahata, T. (2024). Plasminogen activator inhibitor-1 promotes immune evasion in tumors by facilitating the expression of programmed cell death-ligand 1. Front Immunol 15, 1365894. 10.3389/fimmu.2024.1365894.

35. Brune, J.E., Chang, M.Y., Altemeier, W.A., and Frevert, C.W. (2021). Type I Interferon Signaling Increases Versican Expression and Synthesis in Lung Stromal Cells During Influenza Infection. J Histochem Cytochem 69, 691–709. 10.1369/00221554211054447.

36. Chang, M.Y., Tanino, Y., Vidova, V., Kinsella, M.G., Chan, C.K., Johnson, P.Y., Wight, T.N., and Frevert, C.W. (2014). A rapid increase in macrophage-derived versican and hyaluronan in infectious lung disease. Matrix Biol 34, 1–12. 10.1016/j.matbio.2014.01.011.

37. Yaseen, M.M., Abuharfeil, N.M., Darmani, H., and Daoud, A. (2020). Mechanisms of immune suppression by myeloid-derived suppressor cells: the role of interleukin-10 as a key immunoregulatory cytokine. Open Biol 10, 200111. 10.1098/rsob.200111.

38. Wang, J.C., and Sun, L. (2022). PD-1/PD-L1, MDSC Pathways, and Checkpoint Inhibitor Therapy in Ph(-) Myeloproliferative Neoplasm: A Review. Int J Mol Sci 23. 10.3390/ijms23105837.

39. Jayakumar, A., and Bothwell, A.L.M. (2019). Functional Diversity of Myeloid-Derived Suppressor Cells: The Multitasking Hydra of Cancer. J Immunol 203, 1095–1103. 10.4049/jimmunol.1900500.

40. Groth, C., Hu, X., Weber, R., Fleming, V., Altevogt, P., Utikal, J., and Umansky, V. (2019). Immunosuppression mediated by myeloid-derived suppressor cells (MDSCs) during tumour progression. Br J Cancer 120, 16–25. 10.1038/s41416-018-0333-1.

41. Jauch-Speer, S.L., Herrera-Rivero, M., Ludwig, N., Veras De Carvalho, B.C., Martens, L., Wolf, J., Imam Chasan, A., Witten, A., Markus, B., Schieffer, B., et al. (2022). C/EBPdelta-induced epigenetic changes control the dynamic gene transcription of S100a8 and S100a9. Elife 11. 10.7554/eLife.75594.

42. Pappritz, K., Voss, I., El-Shafeey, M., and Van Linthout, S. (2025). Sex differences in age-related cardiac and splenic S100A9 and NLRP3 expression. J Leukoc Biol. 10.1093/jleuko/qiaf031.

43. Stadtmann, A., and Zarbock, A. (2012). CXCR2: From Bench to Bedside. Front Immunol 3, 263. 10.3389/fimmu.2012.00263.

44. Li, S., Feng, J., Wu, F., Cai, J., Zhang, X., Wang, H., Fetahu, I.S., Iwanicki, I., Ma, D., Hu, T., et al. (2020). TET2 promotes anti-tumor immunity by governing G-MDSCs and CD8(+) T-cell numbers. EMBO Rep 21, e49425. 10.15252/embr.201949425.

45. Cai, Z., Kotzin, J.J., Ramdas, B., Chen, S., Nelanuthala, S., Palam, L.R., Pandey, R., Mali, R.S., Liu, Y., Kelley, M.R., et al. (2018). Inhibition of Inflammatory Signaling in Tet2 Mutant Preleukemic Cells Mitigates Stress-Induced Abnormalities and Clonal Hematopoiesis. Cell Stem Cell 23, 833–849 e835. 10.1016/j.stem.2018.10.013.

46. Barrios, E.L., Rincon, J.C., Willis, M., Polcz, V.E., Leary, J.R., Darden, D.B., Balch, J.A., Larson, S.D., Loftus, T.J., Mohr, A.M., et al. (2024). Transcriptomic Differences in Peripheral Monocyte Populations in Septic Patients Based on Outcome. Shock 62, 208–216. 10.1097/SHK.0000000000002379.

47. Balch, J.A., Chen, U.I., Liesenfeld, O., Starostik, P., Loftus, T.J., Efron, P.A., Brakenridge, S.C., Sweeney, T.E., and Moldawer, L.L. (2023). Defining critical illness using immunological endotypes in patients with and without sepsis: a cohort study. Crit Care 27, 292. 10.1186/s13054-023-04571-x.

48. Sherwood, E.R., Burelbach, K.R., McBride, M.A., Stothers, C.L., Owen, A.M., Hernandez, A., Patil, N.K., Williams, D.L., and Bohannon, J.K. (2022). Innate Immune Memory and the Host Response to Infection. J Immunol 208, 785–792. 10.4049/jimmunol.2101058.

49. Rincon, J.C., Wang, D., Polcz, V.E., Barrios, E.L., Dirain, M.L., Ungaro, R.F., Nacionales, D.C., Zeumer-Spataro, L., Xiao, F., Efron, P.A., et al. (2025). Innate immune training in the neonatal response to sepsis. Molecular medicine 31, 159. 10.1186/s10020-025-01179-5.

50. Nacionales, D.C., Gentile, L.F., Vanzant, E., Lopez, M.C., Cuenca, A., Cuenca, A.G., Ungaro, R., Li, Y., Baslanti, T.O., Bihorac, A., et al. (2014). Aged mice are unable to mount an effective myeloid response to sepsis. J Immunol 192, 612–622. 10.4049/jimmunol.1302109.

51. Bible, L.E., Pasupuleti, L.V., Gore, A.V., Sifri, Z.C., Kannan, K.B., and Mohr, A.M. (2015). Chronic restraint stress after injury and shock is associated with persistent anemia despite prolonged elevation in erythropoietin levels. J Trauma Acute Care Surg 79, 91–96; discussion 96–97. 10.1097/TA.0000000000000686.

52. JR, S., MA, H., T, S., Y, K., CE, P., NH, N., RP, D., R, P., K, I., S, H., et al. (10/26/2015). High Fractional Occupancy of a Tandem Maf Recognition Element and Its Role in Long-Range β-Globin Gene Regulation - PubMed. Molecular and cellular biology 36. 10.1128/MCB.00723-15.

53. P, K., ML, G., CE, P., RP, D., K, K., H, B., E, M., A, R., MP, K., and R, B. (12/11/2021). Methylscaper: an R/Shiny app for joint visualization of DNA methylation and nucleosome occupancy in single-molecule and single-cell data - PubMed. Bioinformatics (Oxford, England) 37. 10.1093/bioinformatics/btab438.

