## Supplemental Material for "Sepsis Induces Age- and Sex-Specific Chromatin Remodeling in Myeloid-Derived Suppressor Cells"

Supplemental Text and Figures
Click here to access/download;Supplemental Text and Figures;Charles_et_al_Supplemental.pdf

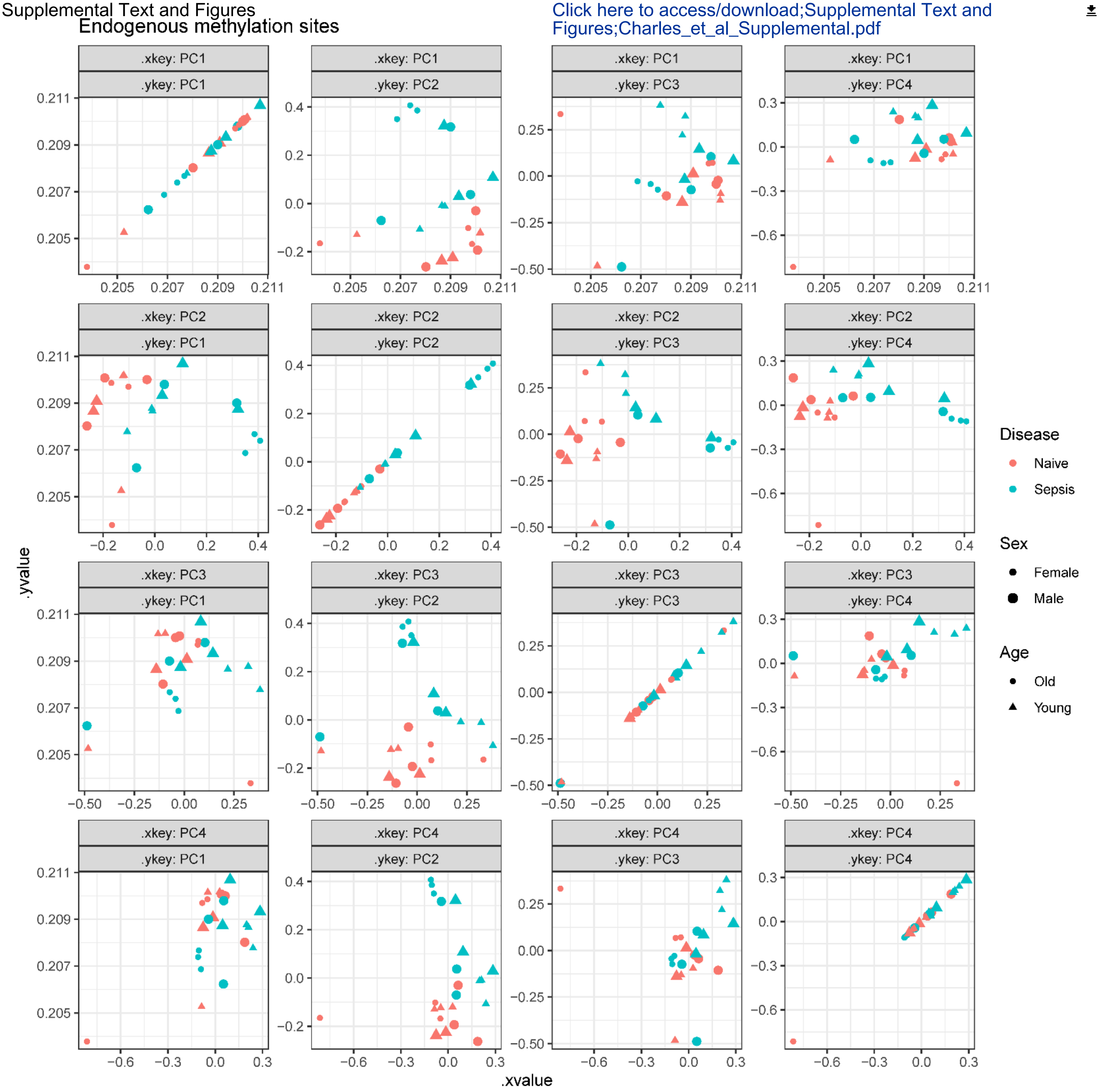


Supplemental Fig 1: **Principal component analysis (PCA) of endogenous CpG methylation (HCG) in splenic MDSCs.** PCA was performed on the percent methylation of CpG (HCG) sites across all covered promoters in CD11b$^{+}$ Gr1$^{+}$ MDSCs from male and female mice, young and old, with or without sepsis. Each point represents a single mouse. Color indicates disease condition (blue = naïve, red = sepsis), shape denotes sex (circle = female, triangle = male), and shading differentiates age (light = young, dark = old). Principal components 1-4 are plotted in pairwise combinations to visualize clustering. Old septic females exhibit distinct separation, indicating strong methylome remodeling in response to sepsis.

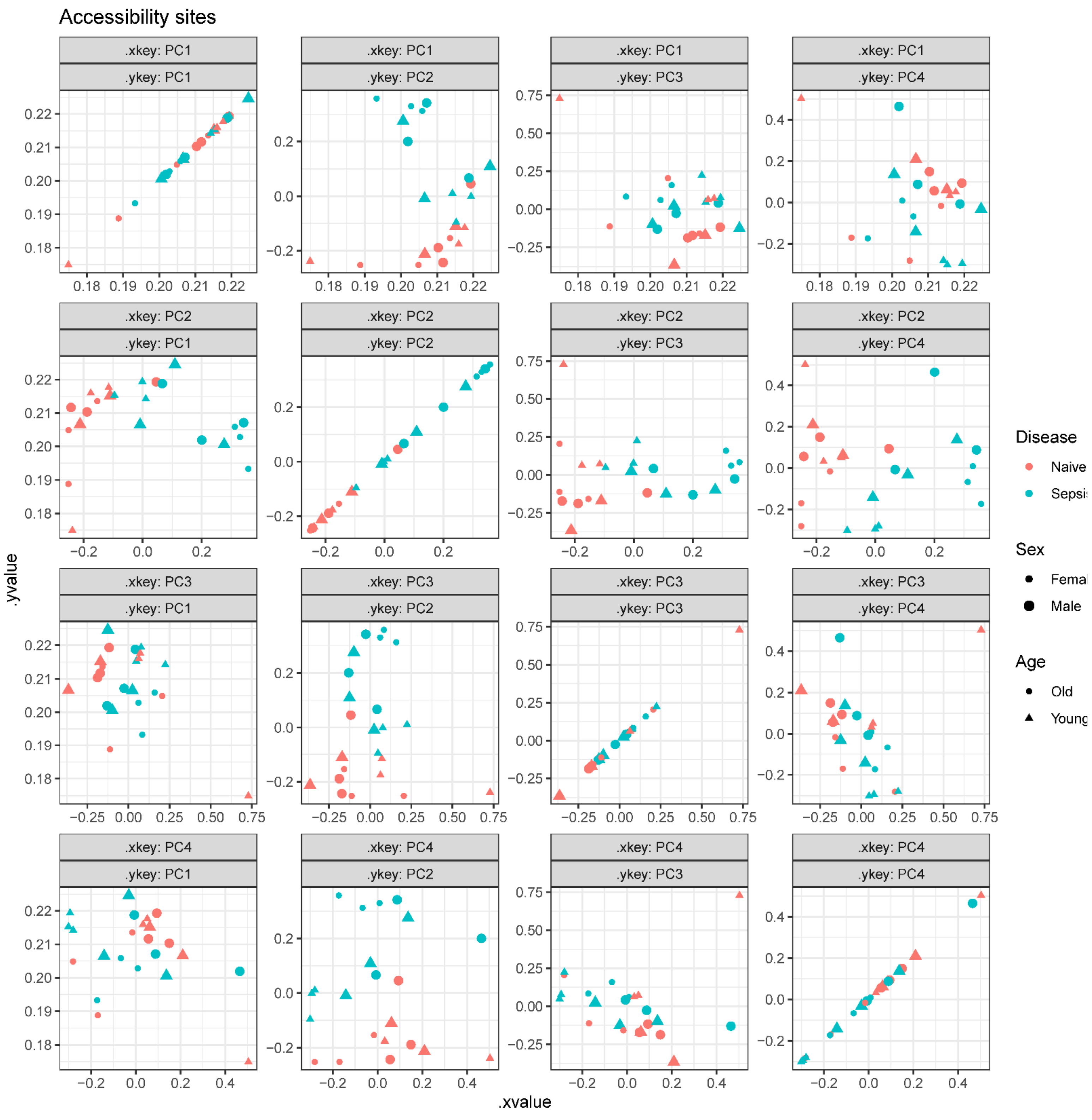


Supplemental Fig 2: **Principal component analysis (PCA) of GpC methylation (chromatin accessibility) in splenic MDSCs.** PCA was performed using GCH methylation values from MAPit-FENGC to reflect promoter accessibility across MDSC samples. Each point represents a single mouse. Color indicates disease condition (blue = naïve, red = sepsis), shape denotes sex (circle = female, triangle = male), and shading differentiates age (light = young, dark = old). Separation by sepsis status is evident across multiple PCs, with the strongest divergence observed in female mice. These results suggest widespread chromatin remodeling following sepsis, particularly in females.

**Supplemental Table 1**

**Gene and Primer List used for MAPit-FENGC**

Adgre1:ENSMUST00000086763
Akt1:ENSMUST00000001780
Ambp:ENSMUST00000030041
Arg1:ENSMUST00000020161
Atf6:ENSMUST00000027974
C3ar1:ENSMUST00000042081
Car4:ENSMUST00000103194
Car6:ENSMUST00000030817
Ccl12:ENSMUST00000000194
Ccl17:ENSMUST00000034232
Ccl2:ENSMUST00000000193
Ccl3:ENSMUST00000001008
Ccl5:ENSMUST00000035938
Ccl7:ENSMUST00000021011
Ccr1:ENSMUST00000026911
Cd247:ENSMUST00000161971
Cd274:ENSMUST00000016640
Cd63:ENSMUST00000219317
Cd74:ENSMUST00000167610
Cd9:ENSMUST00000032492
Cebpb:ENSMUST00000070642
Ctsl:ENSMUST00000222517
Cxcl2:ENSMUST00000200681
Cxcl3:ENSMUST00000031326
Cxcr2:ENSMUST00000106899
Cybb:ENSMUST00000015484
Dab2:ENSMUST00000159552
Dcn:ENSMUST00000105287
Dnmt3b:ENSMUST00000072997
Egln3:ENSMUST00000039516
Emp1:ENSMUST00000205156
F7:ENSMUST00000033820
Fabp7:ENSMUST00000020024
Fam20c:ENSMUST00000026972
Fgr:ENSMUST00000030693
Fosb:ENSMUST00000003640
Fyb:ENSMUST00000090461

Gpnmb:ENSMUST00000204260
Gprc5b:ENSMUST00000008878
Hdac3:ENSMUST00000043498
Htr2a:ENSMUST00000036653
Ighm:ENSMUST00000177715
Igkv12-44:ENSMUST00000103367
Igkv12-46:ENSMUST00000103365
Igkv4-69:ENSMUST00000103349
Igkv8-24:ENSMUST00000103384
Igkv9-120:ENSMUST00000103316
Iglv1:ENSMUST00000103746
Il10:ENSMUST00000016673
Il1r2:ENSMUST00000027243
Il1rl2:ENSMUST00000194296
Il4ra:ENSMUST00000033004
Il6:ENSMUST00000026845
Irf8:ENSMUST00000160943
Jun:ENSMUST00000107094
Kdm6b:ENSMUST00000094077
Lcn2:ENSMUST00000050785
Lgals1:ENSMUST00000089377
Lgals2:ENSMUST00000044584
Lgals3:ENSMUST00000151405
Lgals9:ENSMUST00000108268
Lrp1:ENSMUST00000049149
Lyz1:ENSMUST00000092162
Mmp19:ENSMUST00000026411
Mmp8:ENSMUST00000018765
Mmp9:ENSMUST00000137626
Mt2:ENSMUST00000034214
Nfkbiz:ENSMUST00000036273
Nos2:ENSMUST00000018610
Nrf1:ENSMUST00000115212
Plac8:ENSMUST00000031264
Pmp22:ENSMUST00000108702
Ptgs2:ENSMUST00000035065
Retn:ENSMUST00000012849
Rnase2a:ENSMUST00000061936
Ros1:ENSMUST00000020045
S100a10:ENSMUST00000045756
S100a8:ENSMUST00000069927

S100a9:ENSMUST00000117167
Saa3:ENSMUST00000006956
Serpina1a:ENSMUST00000072876
Serpinb1a:ENSMUST00000076352
Serpine1:ENSMUST00000041388
Slc7a2:ENSMUST00000057784
Sparc:ENSMUST00000214685
Spp1:ENSMUST00000086833
Stab1:ENSMUST00000036618
Stat3:ENSMUST00000103114
Tet2:ENSMUST00000098603
Tgfb1:ENSMUST00000002678
Timp3:ENSMUST00000020234
Tmem176a:ENSMUST00000204482
Tmem176a:ENSMUST00000204482
Tnfaip3:ENSMUST00000019997
Vcan:ENSMUST00000109546
Vdr:ENSMUST00000023119
Vegfa:ENSMUST00000071648
Vnn1:ENSMUST00000041416
Vsig4:ENSMUST00000050707
Atp6v0d2:ENSMUST00000029900
Dnase1l3:ENSMUST00000026315
Gm5849:ENSMUST00000180151
Hdac9-1
Hdac9-2
Igkv1-117:ENSMUST00000103317
Igkv6-15:ENSMUST00000103393
Igkv8-28:ENSMUST00000197525
Trem1:ENSMUST00000048782
Map3k1:ENSMUST00000109267
Ikbkg:ENSMUST00000164101
Ash1l:ENSMUST00000186583
Btk:ENSMUST00000033617
Cd180:ENSMUST00000170878
Cxcl15:ENSMUST00000031322
Cxcr4:ENSMUST00000052172
Foxo1:ENSMUST00000053764
Gdf5:ENSMUST00000040162
Hsf1:ENSMUST00000226860
Hsf2bp:ENSMUST00000238192

Hsf4:ENSMUST00000173102
Hsf5:ENSMUST00000093956
Hsfy2:ENSMUST00000062085
Hsp90aa1:ENSMUST00000155242
Hsp90ab1:ENSMUST00000024739
Hspa1a:ENSMUST00000087328
Hspa1b:ENSMUST00000172753
Hspa2:ENSMUST00000080449
Ifna1:ENSMUST00000094972
Ifna2:ENSMUST00000105147
Ifng:ENSMUST00000068592
Ikbkb:ENSMUST00000033939
Il12a:ENSMUST00000029345
Il12b:ENSMUST00000102796
Il3ra:ENSMUST00000224163
Il6:ENSMUST00000026845
Irak1:ENSMUST00000114354
Irak4:ENSMUST00000109248
Irf3:ENSMUST00000003284
Irf5:ENSMUST00000167252
Irf7:ENSMUST00000106023
Lbp:ENSMUST00000146600
Ly86:ENSMUST00000021860
Mapk1:ENSMUST00000232611
Mapk13:ENSMUST00000233984
Mapk14:ENSMUST00000004990
Mapk15:ENSMUST00000160092
Mapk3:ENSMUST00000050201
Mapk4:ENSMUST00000091851
Mapk6:ENSMUST00000049355
Mapk8:ENSMUST00000111945
Mme:ENSMUST00000194134
Myd88:ENSMUST00000035092
Nfkb1:ENSMUST00000029812
Nfkb2:ENSMUST00000237330
Nfkbia:ENSMUST00000021413
Nfkbib:ENSMUST00000032815
Nr2c2:ENSMUST00000113460
Pik3ap1:ENSMUST00000059672
Pik3c2a:ENSMUST00000170430
Pik3c2g:ENSMUST00000218528

Pik3c3:ENSMUST00000115812
Pik3ca:ENSMUST00000108243
Pik3r1:ENSMUST00000055518
Pik3r2:ENSMUST00000034296
Pik3r3:ENSMUST00000030464
Pik3r5:ENSMUST00000021283
Pik3r6:ENSMUST00000102613
Ptpn6:ENSMUST00000171549
Rela:ENSMUST00000025867
Src:ENSMUST00000109533
Tab1:ENSMUST00000229320
Tab2:ENSMUST00000146444
Tlr2:ENSMUST00000029623
Tlr4:ENSMUST00000107365
Tnf:ENSMUST00000025263
Traf6:ENSMUST00000004949
Vav1:ENSMUST00000169220
